# Spatiotemporal Modulations in Heterotypic Condensates of Prion and α-Synuclein Control Phase Transitions and Amyloid Conversion

**DOI:** 10.1101/2021.09.05.459019

**Authors:** Aishwarya Agarwal, Lisha Arora, Sandeep K. Rai, Anamika Avni, Samrat Mukhopadhyay

## Abstract

Biomolecular condensates formed via liquid-liquid phase separation (LLPS) of proteins and nucleic acids are thought to govern critical cellular functions. These multicomponent assemblies provide dynamic hubs for competitive homotypic and heterotypic interactions. Here, we demonstrate that the complex coacervation between the prion protein (PrP) and α-synuclein (α-Syn) within a narrow stoichiometry regime results in the formation of highly dynamic liquid droplets. Domain-specific electrostatic interactions between the positively charged intrinsically disordered N-terminal segment of PrP and the negatively charged C-terminal domain of α-Syn drive the formation of these highly tunable, reversible, thermo-responsive condensates. Picosecond time-resolved measurements revealed the existence of relatively ordered electrostatic nanoclusters that are stable on the nanosecond timescale and can undergo breaking-and-making on a much slower timescale giving rise to the liquid-like behavior on the second timescale and mesoscopic length-scale. The addition of RNA to these preformed coacervates yields multiphasic, anisotropic, vesicle-like, hollow condensates. LLPS promotes liquid-to-solid maturation of α-Syn-PrP condensates resulting in the rapid conversion into heterotypic amyloids. Our results suggest that synergistic interactions between PrP and α-Syn in liquid condensates can offer mechanistic underpinnings of their physiological role and overlapping neuropathological features.

## Introduction

The precise spatiotemporal regulation of cellular machinery in its naturally crowded milieu is critical for the sustenance of life. Emerging evidence suggests a central role of liquid-liquid phase separation (LLPS) in maintaining subcellular organization via the formation of highly dynamic protein and nucleic-acid-rich biomolecular condensates, also known as membrane-less organelles.^1–9^ Due to the absence of any delimiting membrane, these intracellular emulsions allow rapid exchange of components within the cellular environment and exhibit liquid-like behavior. These assemblies are highly context-dependent and facilitate an array of complex cellular functions ranging from chromatin reorganization to transcriptional regulation. Given the multicomponent nature and complex functions associated with these condensates, they display different internal architectures.^10–12^ For instance, nucleoli display distinct nested sub-compartments and hierarchal organization.^13^ The components inside these condensates have been broadly classified into two categories namely, scaffolds and clients.^14,15^ Scaffolds refer to the resident biomolecules with multiple interaction motifs which drive the formation of these multi-component condensates via a dense network of intermolecular contacts involving electrostatic, hydrophobic, hydrogen bonding, dipole-dipole, π-π, and cation-π interactions.^16–19^ On the other hand, biomolecules recruited via direct interaction with scaffolds, which are otherwise not required for the condensate formation are referred to as clients.^9,15^ The inherently multivalent intrinsically disordered proteins/regions (IDPs/IDRs) with prion-like domains and low-complexity regions are the most common scaffolds governing the formation of these percolated coacervates.^20–28^ The physical origin of these condensates is dictated by the sequence architecture and composition of the scaffolds.^27,29,30^ These assemblies are often concentrated in putative RNA binding proteins such as Fused in Sarcoma (FUS) and FUS family proteins such as transactive response DNA binding protein 43 (TDP-43), hnRNPA1, and so forth.^4^ Therefore, aberrant phase transitions of these protein-rich droplets can also promote deleterious fibrillization that is associated with various neurodegenerative diseases including Alzheimer’s disease, amyotrophic lateral sclerosis, frontotemporal dementia, and so forth.^31–36^

In recent years, the emergence of overlapping neuropathological features has raised the possibility of heterologous aggregation of different amyloidogenic proteins. In general, the co-existence of distinct pathologies is attributed to the synergistic interaction between different proteins, cross-seeding between unrelated proteins, or receptor-mediated toxicity.^37^ For instance, a wealth of evidence suggests proximal locations or colocalizations of aggregates of unrelated amyloidogenic proteins such as α-synuclein (α-Syn), tau, amyloid-β, TDP-43 in the brains of patients.^38–41^ Along the same line, abnormal deposits of α-Syn in the form of Lewy bodies have been found in patients with sporadic or genetic prion diseases such as Creutzfeldt-Jakob disease (CJD) which is linked to the misfolding of the prion protein (PrP).^42^Although PrP and α-Syn are independently known to aggregate and form cytoplasmic inclusions, their coexistence raises important questions about the underlying molecular mechanism. Recent studies have indicated that PrP acts as a receptor for α-syn oligomers and fibrils triggering the downstream signaling cascade resulting in cellular toxicity.^43–46^ Also, inoculation of infectious prions in aged α-Syn transgenic mice has been associated with extensive α-Syn deposits.^47^

α-Syn is a 140-residue neuronal IDP, aggregation of which is associated with Parkinson’s disease. α-Syn contains three distinct regions namely, an amphipathic lysine-rich amino terminus (residues 1-60) with a highly conserved lipid-binding region, a central hydrophobic region known as the non-amyloid-β-component (NAC; residues 61-95) essential for aggregation, and an acidic carboxy-tail (residues 96-140) that interacts with metal ions and other proteins (Fig. 1a, b).^48^ Whereas PrP is a 253-residue C-terminally glycophosphatidylinositol (GPI)-anchored protein consisting of two distinct regions namely, a highly flexible intrinsically disordered N-terminal segment (residues 23-120) and a globular C-terminal domain (residues 121-231) (Fig. 1c, d).^49,50^ The N-terminal IDR harbors several key regions such as the polybasic region comprising two lysine clusters (residues 23-30 and 100-110), a glycine-rich octapeptide repeat region (residues 51-90), and a hydrophobic segment (113-135).^51,52^ The globular C-terminal domain of PrP consists of three α-helices and two short antiparallel β-strands. The misfolding and aggregation of PrP has been associated with a class of invariably fatal and transmissible neurodegenerative diseases such as CJD. In order to elucidate the molecular basis of their overlapping neuropathology, we set out to study the interaction between human α-Syn and PrP. In this work, we demonstrate that phase separation via a complex coacervation of α-Syn and PrP yields highly dynamic heterotypic condensates comprising ephemeral electrostatic nanoclusters within the liquid-like mesoscopic organization. Domain-specific interactions and charge anisotropies provide spatiotemporal modulations of these highly tunable and thermo-responsive condensates that eventually undergo maturation into highly ordered, heterotypic, solid-like amyloid fibrils.

**Fig. 1.**
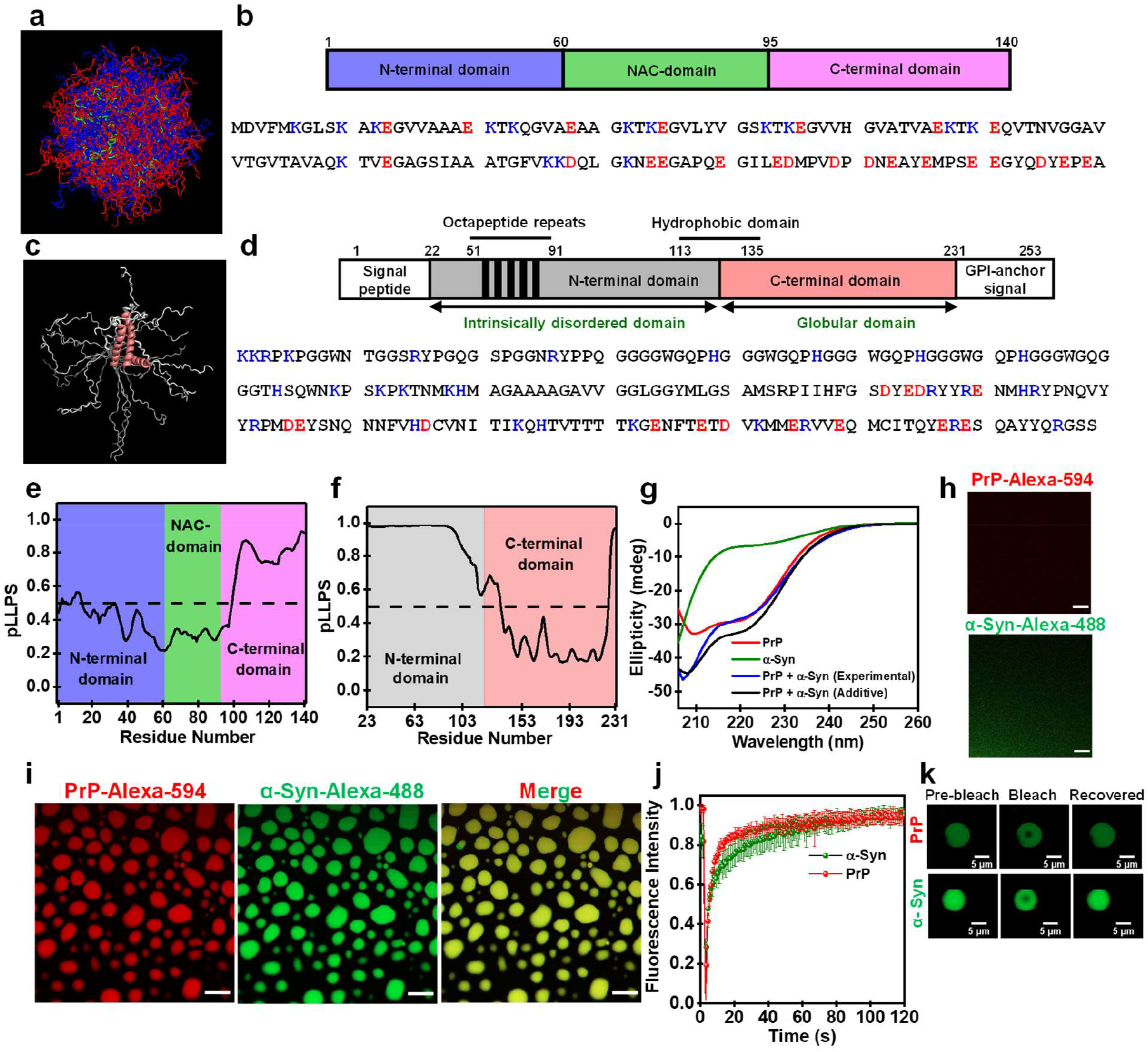
Heterotypic phase separation of α-Syn and PrP. **a** An overlay of 576 conformations obtained from the ensemble structure of α-Syn (PED ID: PED00024e001 generated using PyMOL (Schrödinger, LLC, New York). **b** Domain architecture and the amino acid sequence of α-Syn. Positively charged negatively charged amino acids are shown in blue and red, respectively. **c** An overlay of 20 conformations obtained from the NMR structure of human PrP (90-231) (PDB ID: 2LSB) generated using PyMOL. **d** Schematic representation of PrP (23-231) indicating the N-terminal disordered and the C-terminal globular domains. Positively charged negatively charged amino acids are shown in blue and red, respectively. Prediction of the LLPS propensity using FuzDrop for **e** α-Syn and **f** PrP (23-231). **g** CD spectra for PrP, α-Syn, PrP + α-Syn upon LLPS (experimentally observed), PrP + α-Syn upon LLPS (additive). **h**,**i** Confocal images of mixed homogeneous phases of PrP and α-Syn and complex coacervates of PrP and α-Syn performed using Alexa-594-labeled PrP (Cys 31) and Alexa-488-labeled α-Syn (Cys 90) indicating their complete miscibility and colocalization within droplets. Scale bar: 10 µm. **j** FRAP kinetics of multiple droplets (∼1% Alexa-488-labeled protein; n = 5) for PrP (red) and α-Syn (olive). **k** Fluorescence images of droplets during FRAP measurements. PrP and α-Syn concentrations were 20 µM and 30 µM, respectively. See Methods for details.

## Results

### Heterotypic phase separation of α-Syn and PrP

In order to make predictions about the phase behavior of PrP and α-Syn, we first set out to evaluate the disorder and charge distribution in the primary amino acid sequence. As expected, disorder predictors revealed significant disorder for the entire sequence of the α-Syn and the N-terminal region of the PrP (Fig. S1a, b). The primary sequence of PrP and α-Syn carries a net positive (∼ +10) and negative charge (∼ -8), respectively, at a near-neutral pH. The linear net charge per residue (NCPR) plots generated using Classification of Intrinsically Disordered Ensemble Regions (CIDER)^53^ showed clustering of positive charges at the N-terminal part of PrP and negative charges preferentially located at the C-terminal part of α-Syn (Fig. S1c, d). We further evaluated the sequence-dependent phase behavior using well-known LLPS predictors FuzDrop^54^ and catGRANULE^55^, which revealed a significant phase separation propensity for both the proteins (Fig. 1e, f, Fig. S1e). It is interesting to note that the phase separation propensity is higher for the positively charged N-terminal segment of PrP and the highly negatively charged C-terminal part of α-Syn. Therefore, we postulated that the electrostatic interactions could potentially promote their complex coacervation at the physiological pH. In order to experimentally verify if these two proteins together can undergo complex coacervation, we began by characterizing their phase behavior *in vitro*. Under our experimental condition, two separate solutions of PrP and α-Syn remained clear and dispersed at near-neutral pH (pH 6.8-7.4, 37 °C). To establish their monomeric nature, we also performed dynamic light scattering (DLS) measurements that revealed a hydrodynamic diameter of ∼ 6 nm and ∼ 10 nm for α-Syn and PrP, respectively as expected for their monomeric hydrodynamic size (Fig. S1f, g).^56,57^ We next co-incubated PrP and α-Syn at different stoichiometries based on their net charges and observed their phase behavior using turbidity measurements and microscopic investigations. Upon the addition of PrP to α-Syn (molar ratio: α-Syn:PrP =1.5, 37 °C), the solution spontaneously turned turbid, indicating the presence of micron-sized condensates as also determined using DLS (Fig. S1h). Microscopic studies showed the presence of spherical liquid-like droplets that undergo fusion. The SDS-PAGE analysis also revealed the presence of both α-Syn and PrP in the sedimented condensed phase, establishing their heterotypic nature (Fig. S1i). Additionally, circular dichroism spectroscopy (CD) revealed no changes in the secondary structural content upon LLPS, indicating that the intrinsic disorder and structural elements are retained within these complex coacervates (Fig. 1g). Together, these experiments indicated the formation of heterotypic liquid-like complex coacervates of α-Syn and PrP.

In order to directly observe the presence of both α-Syn and PrP in these liquid droplets, we performed two-color confocal fluorescence imaging. We created single cysteine mutants at residues 90 and 31 of α-Syn and PrP, respectively, to perform site-specific fluorescence labeling using thiol-active fluorescent dyes namely, AlexaFluor-488 (green) and AlexaFluor-594 (red). PrP and α-Syn doped with their respective fluorescently labeled proteins (∼1%) were mixed and observed using confocal microscopy, which revealed their complete colocalization within these liquid droplets (Fig. 1h, i). The condensed phase concentration of α-Syn and PrP in these droplets was estimated to be ∼ 10 mM and ∼ 15 mM, respectively, compared to the dispersed phase concentration of ∼ 15 µM and ∼ 25 µM, respectively (Fig. S1j). We next examined the internal mobility of both these proteins within the condensates by utilizing fluorescence recovery after photobleaching (FRAP) kinetics. Both PrP and α-Syn revealed fast and complete recovery indicating their fast translational diffusion within these condensates (Fig. 1j, k). Taken together, these studies highlight the highly dynamic nature of these heterotypic condensates in which both α-Syn and PrP exhibit complete miscibility and high molecular mobility within the mesoscopic condensed phase. These heterotypic condensates are likely to be formed via electrostatic interactions between oppositely charged disordered domains of the two proteins similar to complex coacervation of polyelectrolytes via charge neutralization that has been previously observed for other proteins and nucleic acids.^58,59^ Therefore, we next set out to unmask the role of the electrostatic effects in heterotypic LLPS of α-Syn and PrP.

### Charge neutralization drives heterotypic LLPS of α-Syn and PrP

Charge neutrality is a primary condition for coacervate formation between counterionic electrolytes which occurs at a well-defined stoichiometry. To this end, we constructed phase diagrams as a function of the protein concentration using turbidity measurements. These measurements in the presence of an increasing concentration of α-Syn at a fixed PrP concentration revealed a typical reentrant phase behavior with three distinct regimes analogous to RNA-induced reentrant phase transitions (Fig. 2a).^11^ LLPS occurred at a narrow stoichiometry regime (molar ratio α-Syn:PrP *≈* 1-4) with the maximum LLPS at α-Syn:PrP molar ratio of *≈*1.5-2. Any deviation from this stoichiometry resulted in the dissolution of these droplets as was also confirmed using fluorescence imaging (Fig. 2b). The dissolution could be due to charge inversion in the presence of excess α-Syn. We next measured the electrophoretic mobility of solution which is indicative of the net surface charge on the peptides. The narrow LLPS regime exhibited an almost neutral net surface charge, whereas higher or lower α-Syn resulted in a charge inversion (Fig. 2c). In addition, a reverse titration experiment with increasing PrP concentration against a fixed α-Syn concentration displayed a similar phase behavior (Fig. S2a). To further support the role of electrostatic interactions, we carried out the phase separation assays as a function of increasing ionic strength. The addition of increasing amounts of salt resulted in the dissolution of these droplets (Fig. 2d). Higher protein concentrations were required to drive LLPS at higher ionic strengths. These results established the critical role of electrostatics in modulating the phase behavior for this counterionic polyelectrolyte mixture. Since electrostatic interactions appear to be the predominant LLPS driver, the next aspect is to investigate the role of temperature in governing the complex coacervation.

**Fig. 2.**
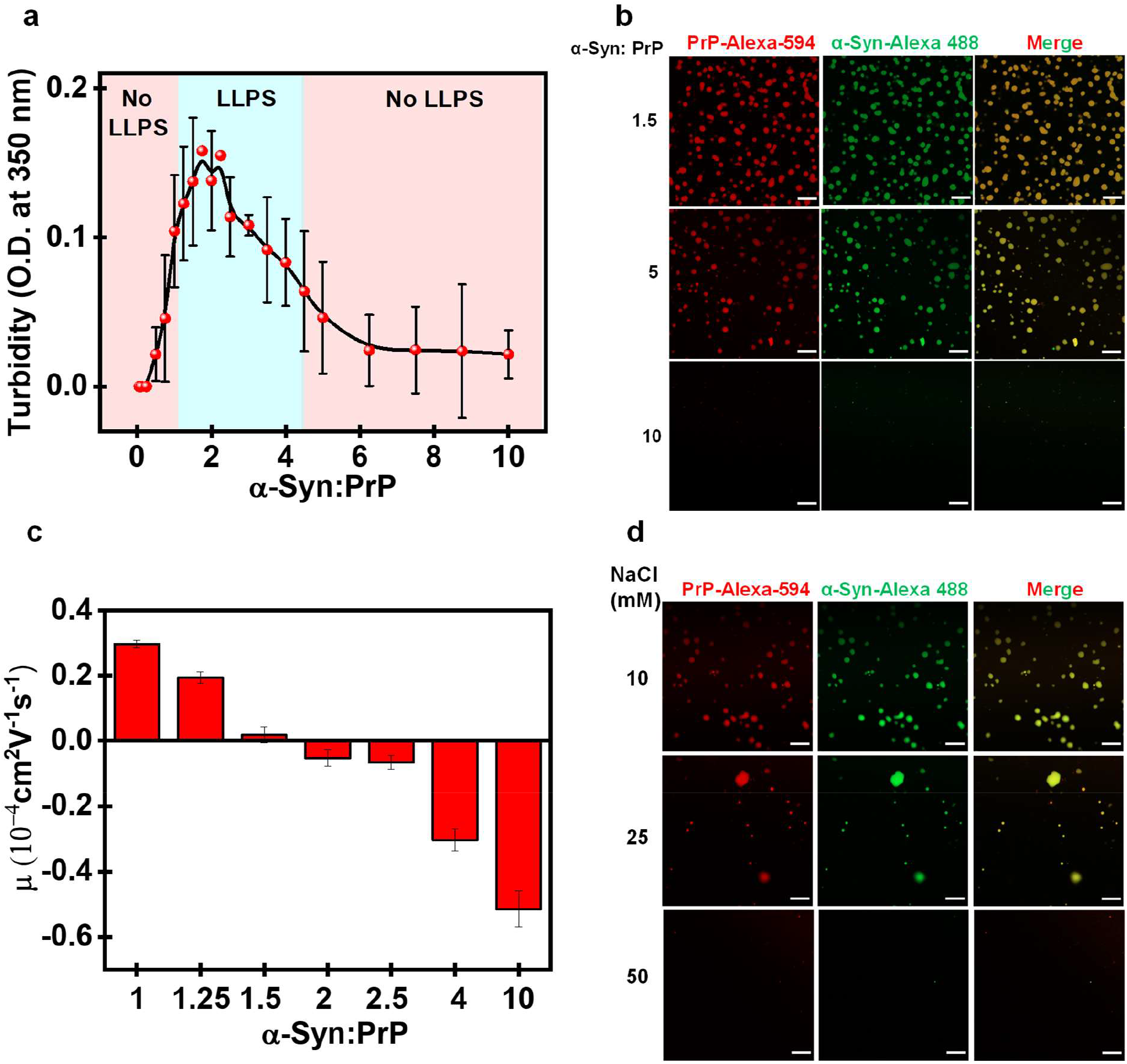
Charge neutralization drives heterotypic LLPS of α-Syn and PrP. **a** Solution turbidity plot at fixed PrP concentration (20 µM) as a function of increasing α-Syn concentrations showing reentrant phase behavior (mean ± SEM; 3). The solid line is for eye guide only. **b** Confocal microscopy images of Alexa-594-labeled PrP and Alexa-488-labeled α-Syn at different stoichiometries as indicated. Scale bar: 10 µm. **c** Electrophoretic mobility (µ) measurements reveal charge inversion with the increase in the α-Syn:PrP ratio. **d** Confocal images of Alexa-594-labeled PrP (20 µM) and Alexa-488-labeled α-Syn (30 µM) droplets with increasing salt concentrations. Scale bar: 10 µm.

We next set out to study the thermo-responsive behavior associated with the complex coacervation of α-Syn and PrP. The phase behavior of a protein is encoded in its amino acid composition and is governed by a critical balance between protein-protein and protein-solvent interactions. The interaction network between protein and solvent is stabilized by contribution from the enthalpy and entropy of mixing which drives the system to attain a minimum global free energy. The tendency of the protein to phase separate may increase or decrease with temperature giving rise to an LCST (lower critical solution temperature) or UCST (upper critical solution temperature) transitions, respectively.^23^ Therefore, we next performed the LLPS assay for these coacervates as a function of temperature ranging from 4 °C to 42 °C. We observed an increase in phase separation propensity as a function of temperature indicating an LCST behavior, which is the signature of an entropy-driven system (Fig. S2b). The entropic gain associated with the release of counterions results in the LCST behavior.^58,60^ The phase behavior was highly tunable as these droplets exhibited temperature-dependent reversibility. Taken together, these results suggest a predominant role of electrostatic interactions in driving this complex coacervation. We postulate that the positively charged N-terminal segment of PrP and negatively charged C-terminal domain of α-Syn participate in these intermolecular interactions. Therefore, we next set out to characterize the role of different domains of both PrP and α-Syn in the two-component phase transition.

### Domain-specific heterotypic interactions drive PrP-α-Syn condensation

In order to unveil the putative role of the negatively charged C-terminal domain of α-Syn in heterotypic LLPS, we created two naturally occurring C-terminally truncated variants of α-Syn, namely 133Stop (α-Syn 1-132) and 103Stop (α-Syn 1-102). These truncated variants are found in insoluble disease deposits associated with synucleinopathies.^61^ The truncation lowers the net negative charge, the charge density, and NCPR. We hypothesized that shortening the acidic C-terminal tail would lower the LLPS propensity. In accordance with our expectation, our turbidity assay revealed a much lower propensity of α-Syn 1-132 to undergo heterotypic LLPS with PrP as compared to full-length α-Syn. α-Syn 1-102 containing an even shorter acidic tail did not exhibit LLPS with PrP (Fig. 3a, Fig. S3a). Our results revealed an imperative role of the C-terminal domain of α-Syn and demonstrated the critical role of charge density in modulating the two-component phase transition of α-Syn and PrP. Similarly, to verify the role of different PrP domains in promoting this heterotypic LLPS, we created naturally occurring truncations in PrP associated with prion diseases. We first questioned the role of the PrP globular domain in its complex coacervation with α-Syn. To this end, we created PrP 112-231 that retains the complete globular domain and a much shorter disordered N-terminal tail. PrP 112-231 did not exhibit LLPS even upon prolonged incubation with α-Syn under our experimental conditions (Fig. 3b, Fig. S3b). We next used a naturally occurring PrP deletion mutant PrP 23-144 (Y145Stop) which retains the disordered N-terminal tail of PrP and is devoid of the C-terminal folded domain.^62,63^ Y145Stop exhibited a similar phase separation propensity as compared to full-length PrP in the presence of α-Syn, indicating the necessary and sufficient role of the N-terminal IDR of PrP in two-component LLPS with α-Syn (Fig. 3b, Fig. S3b).

**Fig. 3.**
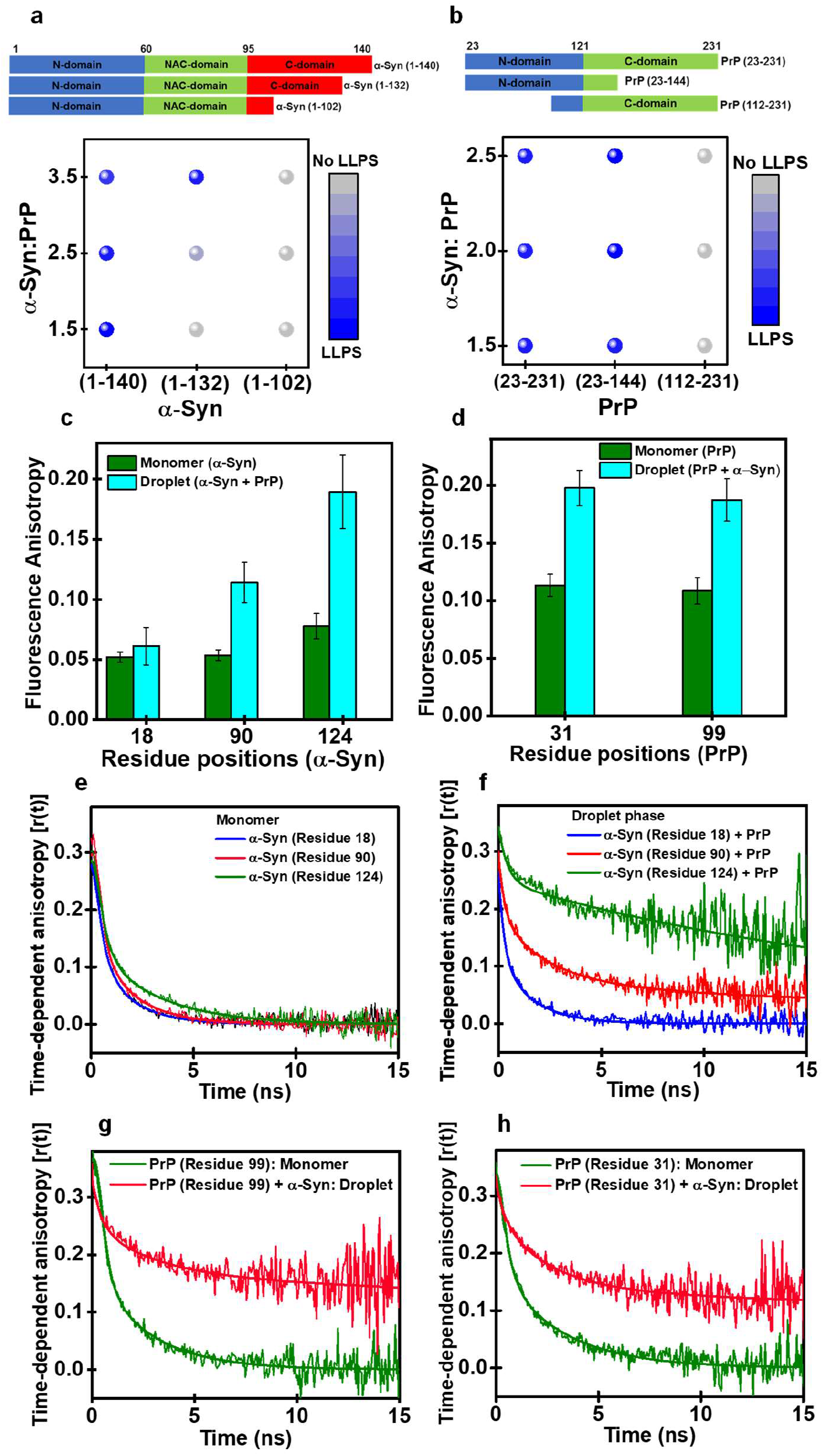
Domain-specific heterotypic interactions and presence of electrostatic clusters within PrP-α-Syn condensates. **a** Schematic representation of different α-Syn constructs used. Phase diagram for different α-Syn constructs at fixed PrP concentration (20 µM) as a function of increasing α-Syn concentrations created from mean turbidity values. **b** Schematic representation of different PrP constructs used. Phase diagram for different PrP constructs (20 µM) as a function of increasing α-Syn concentrations created from mean turbidity values. **c** Steady-state fluorescence anisotropy of single-Cys α-Syn labeled at different positions using F5M in the mixed monomer and droplets (mean ± SEM; 4) **d** Steady-state fluorescence anisotropy of PrP labeled at different positions using F5M in the monomer and droplets (mean ± SEM; 4). **e**,**f** Time-resolved anisotropy decays of F5M-labeled α-Syn in dispersed α-Syn monomers and complex coacervate of PrP-α-Syn. **g**,**h** Time-resolved anisotropy decays of F5M-labeled PrP in dispersed PrP monomer and complex coacervate of PrP-α-Syn. The solid lines are fits obtained using the biexponential and triexponential decay analysis for monomers and droplets, respectively. See Methods, for details of picosecond time-resolved anisotropy decays measurements, analysis, and the estimation of R_h_.

Together, these results showcase the importance of electrostatic effects of two oppositely charged intrinsically disordered domains in complex coacervation. Our experimental observations are in accordance with bioinformatic predictions and demonstrate the critical role of PrP N-terminal and α-Syn C-terminal in dictating the two-component phase behavior. Previous studies with non-biological polyelectrolytes as well as IDPs have revealed the formation of primary units as a preliminary step to the formation of these heterotypic coacervates.^64,65^ Once formed, these primary units coalesce into a highly dynamic liquid-like dense phase that appears to be homogeneous at the mesoscopic length-scale but may contain some molecular/nanoscale order and dynamic heterogeneity. We hypothesized that region-specific electrostatic interactions could potentially induce short length-scale molecular or nanoscale ordering within the condensed phase. Therefore, in order to delineate the short-range order and dynamic heterogeneity at a high spatiotemporal resolution, we next utilized site-specific picosecond time-resolved fluorescence measurements that offer valuable residue-specific dynamic insights on the picosecond to nanosecond timescales.

### Presence of electrostatic nanoclusters within liquid droplets

To address the region-specific structural ordering, we utilized site-specific fluorescence anisotropy measurements that allow us to capture the local rotational flexibility. An increase in the steady-state fluorescence anisotropy at a residue location is attributed to an increase in order and a loss in conformational flexibility. In order to record the region-specific anisotropy, we took advantage of the fact that α-Syn is devoid of Cys and created single Cys mutants at residues 18, 90, and 124 located at the N-terminal domain, NAC domain, and C-terminal domain, respectively, and labeled them using a thiol-active dye namely fluorescein-5-maleimide (F5M). The steady-state fluorescence anisotropy of residue 18 located at the N-terminal domain of α-Syn did not exhibit any changes upon LLPS with PrP, whereas a C-terminal location of α-Syn at residue 124 showed a significant rise in the anisotropy upon phase separation with PrP indicating a critical contribution of the C-terminal domain of α-Syn in LLPS (Fig. 3c). The NAC-domain location α-Syn at residue 90 exhibited only a moderate increase in the anisotropy which could hint at a possible role of hydrophobic residues in promoting such nanoscale ordering (Fig. 3c). To verify if the positively charged N-terminal disordered domain of PrP experiences a similar rotational constraint upon complex coacervation with α-Syn, we created single Cys mutants at residues 31 and 99 of PrP and labeled them using F5M. A similar increase in the fluorescence anisotropy upon LLPS indicated a rotational hindrance for the N-terminal domain of PrP upon LLPS with α-Syn (Fig. 3d). The addition of salt disperses the droplets and results in lower anisotropy values similar to the dispersed phase indicating that the rise of anisotropy was indeed due to local ordering upon the complex coacervation (Fig. S3c). Together these results revealed that the highly basic N-terminal segment of PrP and the acidic C-terminal tail of α-Syn experience restricted rotational mobility within liquid droplets, corroborating our findings that domain-specific heterotypic electrostatic interactions drive the formation of two-component condensates of PrP and α-Syn. However, steady-state fluorescence measurements provide time-averaged information, and therefore, cannot distinguish between various modes of rotational dynamics of polypeptide chains. We next employed picosecond time-resolved fluorescence anisotropy measurements to disentangle distinct dynamical events on the nanosecond timescale.

Picosecond time-resolved fluorescence anisotropy decays yield fluorescence depolarization kinetics that permits us to capture the various modes of rotational relaxation of a polypeptide chain.^66^ For instance, monomeric disordered conformers display a typical rapid depolarization that can be described by biexponential decay kinetics comprising a fast (sub-nanosecond) rotational correlation time representing the local fluorophore dynamics and a slower (nanosecond) rotational correlation time corresponding to the backbone dihedral rotations. For expanded IDPs, the slower correlation time represents a characteristic relaxation time (∼ 1.4 ns) that arises due to short-range torsional fluctuations in the Φ-Ψ dihedral space.^66,67^ As expected, in the monomeric dispersed phase, α-Syn exhibited such typical location-independent biexponential fluorescence depolarization kinetics for all the three locations (residues 18, 90, and 124) that is expected for a rapidly fluctuating expanded disordered state (Fig. 3e). Upon phase separation with PrP, the N-terminal location of α-Syn containing residue 18 retained the dynamic characteristics of a disordered chain, whereas, the C-terminal location at residue 124 exhibited significantly slower depolarization kinetics with a much slower rotational correlation time ∼ 54 ns indicating local clustering around this region of α-Syn (Fig. 3f, Fig. S3d, Table S2). The NAC-domain position (residue 90) showed a moderate increase in the slower rotational correlation time as compared to the dispersed form (Fig. 3f). Similarly, the N-terminal segment of PrP that is highly dynamic in the monomeric dispersed form exhibited significant dampening of rotational dynamics (Fig. 3g, h). These results are in accordance with our hypothesis that electrostatic effects between the acidic C-terminal domain of α-Syn and basic N-terminal segment of PrP are the key drivers of α-Syn-PrP complex coacervation. Such domain-specific electrostatic interactions can yield partially and temporally ordered heterotypic oligomeric domains that can act as non-covalent crosslinks responsible for phase transitions. The hydrodynamic radius (R_h_) of such heterotypic domains estimated from the slower rotational correlation time (∼ 54 ns) by using the well-known Stokes-Einstein relationship is ∼ 4.3 nm (See Table S2).^68^ We would like to note that this is an approximation and the exact hydrodynamic size of the cluster will be dependent on the internal droplet viscosity. Nevertheless, our fluorescence depolarization kinetics indicated electrostatic clustering of C-terminal α-Syn and N-terminal PrP into relatively ordered nano-blobs acting as primary units of liquid droplets. We posit that these clusters that are detected on the nanosecond timescale can undergo breaking-and-making of transient cross-links on a much slower timescale resulting in a liquid-like behavior on a longer length scale as observed by FRAP. Such spatiotemporal regulations might be relevant for aberrant phase transitions and the modulation of the phase behavior by RNA and other biomolecules.

### RNA participates in a competitive multicomponent coacervation

Most phase-separated condensates are enriched in RNA and harbor proteins with RNA binding domains.^4^ Competing protein-protein and protein-RNA interactions underlie the compositional specificity of these cellular condensates in a context-dependent manner.^69,70^ Therefore, we next set out to characterize the role of RNA in regulating this phase transition. PrP contains an RNA-binding domain and is known to interact with nucleic acids.^71,72^ We first tested the phase separation propensity of PrP and α-Syn separately in the presence of RNA. PrP exhibited LLPS in the presence of RNA and displayed reentrant phase behavior (Fig. S4a). However, α-Syn did not undergo phase separation upon the addition of RNA, as reported previously.^31^ We next studied the behavior of preformed PrP-α-Syn complex coacervates in the presence of RNA. Turbidity measurements complemented with microscopic observations revealed a reentrant phase transition of these heterotypic condensates in the presence of RNA under our experimental conditions (Fig. 4a). Low RNA/protein ratios promoted phase separation resulting in the formation of ternary droplets that appear to have a uniform distribution (Fig. 4b). However, beyond a threshold RNA concentration, it resulted in multiphasic droplets with vesicle-like morphology (Fig. 4a, b). Previous studies have also shown the formation of these vesicle-like hollow condensates in the presence of RNA.^11,73^ We hypothesize that the repulsion due to the charge inversion together with a competition between α-Syn and RNA for binding sites on PrP resulted in the formation of these multiphasic condensates beyond a certain RNA concentration. Our electrophoretic mobility measurements supported the charge inversion phenomenon due to a high net negative surface charge at higher RNA concentrations (Fig. 4a inset). The addition of RNA to the preformed PrP-α-Syn coacervates resulted in the displacement of α-Syn from these droplets, indicating a stronger affinity of RNA towards PrP. The displacement of α-Syn by RNA was also confirmed using steady-state fluorescence anisotropy. F5M-labeled residue 124 of α-Syn revealed a sharp increase in the fluorescence anisotropy upon LLPS; however, the addition of increasing concentration of RNA progressively disrupted its interaction with PrP, as revealed by the gradual decrease in the fluorescence anisotropy (Fig. 4c). In addition, SDS-PAGE of sedimented droplets in the presence of RNA indicated a much lower amount of α-Syn, suggesting its weak partitioning into these RNA-rich condensates (Fig. S4 b). To test if these hollow condensates exhibit spatial ordering, we measured fluorescence anisotropy for PrP, which is highly enriched within these vesicle-like condensates. The F5M-labeled residue 31 of PrP exhibited a sharp increase in the anisotropy upon addition of RNA, indicating the presence of molecular ordering within these multiphase condensates (Fig. 4d). Such a molecular ordering might be generic to nucleoprotein vesicles, as has also been shown previously.^73^ Taken together, these results indicate a switch-like behavior of RNA in tuning the compositional specificity of these condensates in a context-dependent manner. It is interesting to note that such interactions can provide spatiotemporal regulations; however, the high enrichment of biomolecules within these condensates makes them a susceptible site for aberrant phase transitions and pathological aggregation under stress conditions. Therefore, we next set out to elucidate the effect of α-Syn-PrP complex coacervation on the aggregation propensity of these heterotypic condensates.

**Fig. 4.**
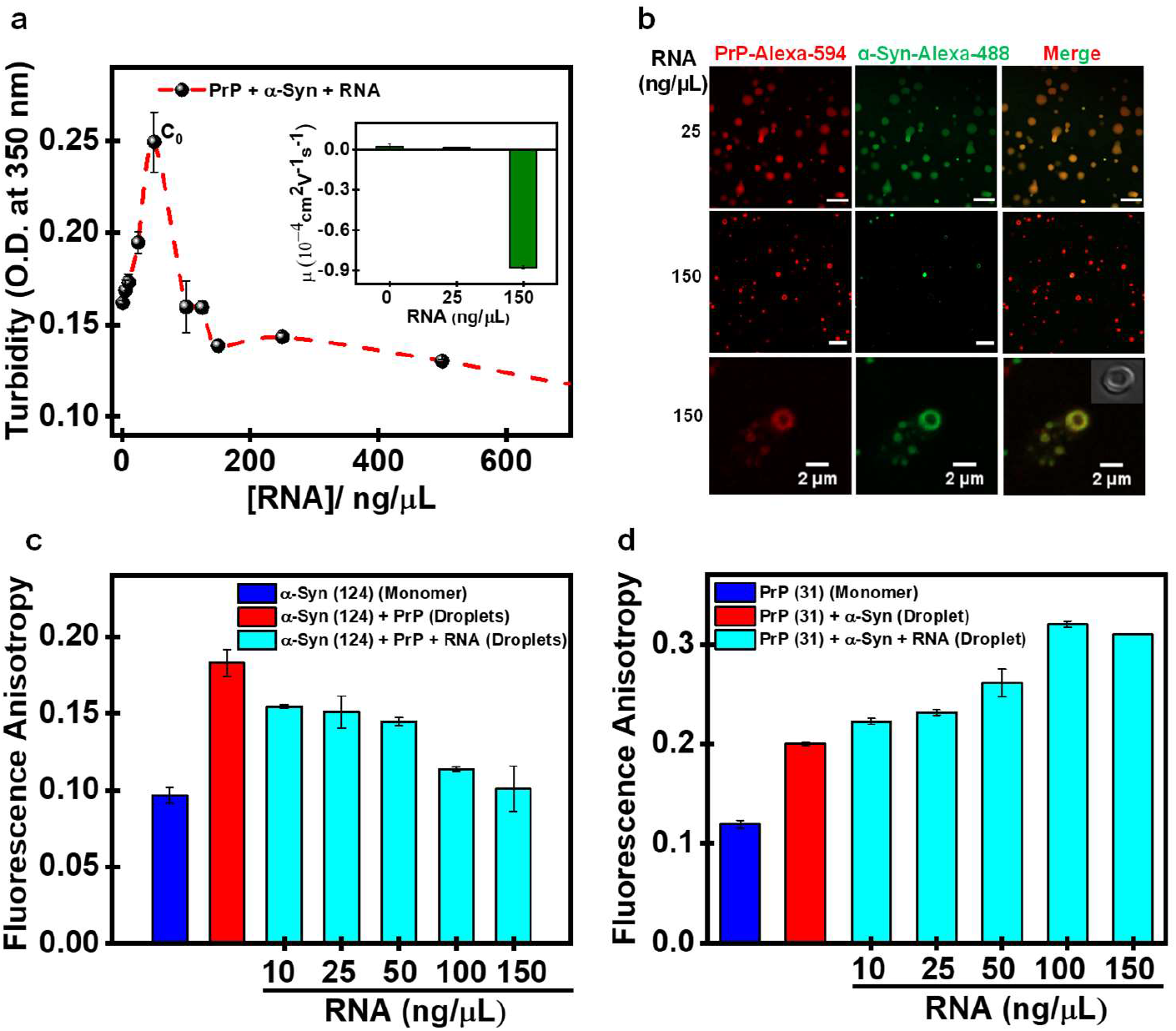
RNA participates in a competitive multicomponent coacervation. **a** Solution turbidity plot as a function of increasing polyU RNA at a fixed α-Syn:PrP ratio showing a reentrant phase behavior. The inset shows charge inversion at different RNA concentrations. **b** Confocal fluorescence images for different regions of the phase diagram with RNA concentrations as indicated. Before the maximum (C_0_), the ternary complex exhibits miscibility, whereas, beyond the maximum (C_0_), the droplets start dispersing by transitioning into multiphasic, vesicle-like, hollow condensates. The inset shows a DIC image for a single hollow condensate. Scale bar: 10 µm **c** Steady-state fluorescence anisotropy for F5M-labeled α-Syn at residue 124 indicating its displacement from the condensates with increasing RNA concentrations. **d** Steady-state fluorescence anisotropy for F5M-labeled PrP residue 31 indicating an increase in the order within hollow condensates with the increase in the RNA concentration.

### Synergistic heterotypic interactions promote liquid-to-solid amyloid transition

LLPS-mediated liquid-to-solid phase transitions into pathological aggregates have been observed for various other neuronal IDPs/IDRs such as FUS, TDP43, tau, and so forth.^74,75^ PrP and α-Syn pathologies have been together implicated in prion diseases. For instance, extensive α-Syn immunoreactive deposits accumulate in the brains of CJD patients.^76^ Also, colocalization of α-Syn and PrP aggregates has been observed in early cytoplasmic inclusions bodies.^42^ Therefore, we next set out to determine if α-Syn-PrP heterotypic interactions within these complex coacervates can promote aggregation and amyloid formation. We monitored the aggregation kinetics of these liquid droplets using a well-known amyloid marker, Thioflavin-T (ThT) (Fig. 5a). α-Syn alone exhibited typical nucleation-dependent polymerization kinetics with a lag phase of ∼ 6 h, as expected, whereas PrP alone did not aggregate under our experimental conditions upon agitation (Fig. 5a). However, heterotypic α-Syn-PrP coacervates upon incubation under a stirring condition rapidly aggregated into amyloids via isodesmic kinetics by completely bypassing the long lag phase (Fig. 5a). In contrast, aggregation at various ratios under non-LLPS conditions did not eliminate the lag phase indicating the critical role of heterotypic condensates in accelerating the amyloid transition via LLPS-mediated pathway (Fig S5a).

**Fig. 5.**
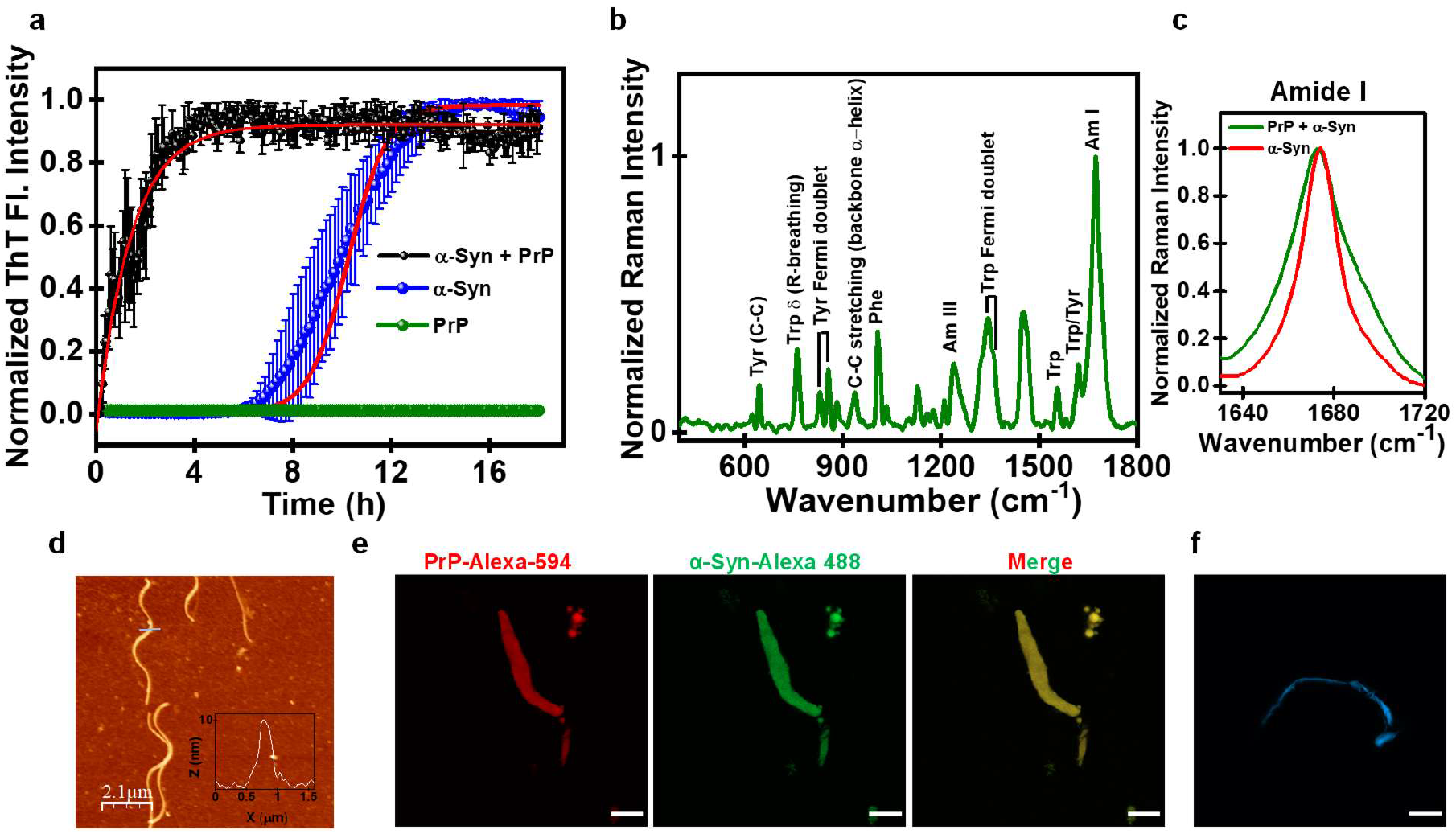
Synergistic heterotypic interactions promote a liquid-to-solid amyloid transition. **a** ThT kinetics for *de novo* aggregation α-Syn (30 µM) and PrP (20 µM) (separately) and LLPS-mediated aggregation via a liquid-to-solid transition of complex coacervates of α-Syn and PrP completely bypassing the lag phase. **b** Vibrational Raman spectra of PrP-α-Syn aggregates indicating their heterotypic nature. **c A**mide I is shown for comparison between PrP-α-Syn heterotypic aggregates formed via LLPS and α-Syn homotypic aggregates formed via *de novo* aggregation. **d** AFM image of LLPS-mediated heterotypic aggregates showing the presence of typical amyloid fibrils. The inset shows the height profile (∼10 nm). **e** Two-color confocal fluorescence images showing colocalization of α-Syn and PrP within these heterotypic aggregates. **f** Confocal fluorescence image of a ThT-positive fibril. Scale bar: 10 µm.

We next structurally characterized these LLPS-mediated heterotypic aggregates using vibrational Raman spectroscopy that allowed us to elucidate the different secondary structural elements present within these aggregates. The amide I vibrational band that originates primarily due to the C=O stretching of the backbone appeared at 1675 cm^-1^, which is the hallmark of a hydrogen-bonded cross-β amyloid architecture (Fig. 5b).^77^ The full-width at half maximum (FWHM) of amide I and III for α-Syn-PrP heterotypic amyloids was considerably broader (31 ± 2 cm^-1^) compared to homotypic α-Syn amyloids (20 ± 1 cm^-1^) (Fig. 5c, Fig. S5c, d). A broader amide I band indicated the contribution from the globular α-helical domain of PrP which, at least in part, is possibly retained in α-Syn-PrP heterotypic amyloids (Fig. 5c). Our CD data also indicated the presence of both helical and β-rich conformers in these amyloids, corroborating our Raman results (Fig. S5b). Additionally, tryptophan residues present in the N-terminal of PrP experience decreased polarity as evident from its small Raman blue-shift to 883 cm^-1^ indicating that the N-terminal part of PrP might be sequestered in the core of α-Syn-PrP amyloids. Further, to visualize these heterotypic amyloid aggregates, we performed atomic force microscopy (AFM) which revealed the presence of typical nanoscopic amyloid fibrils with a height of ∼ 10 nm (Fig. 5d and inset). Additionally, the two-color high-resolution Airy scan confocal fluorescence imaging revealed colocalization of PrP and α-Syn within these amyloid fibrils (Fig. 5e). Together, these results reveal synergistic effects of α-Syn and PrP within liquid-like complex coacervates allowing their conformational sequestration and rapid conversion into highly ordered, solid-like ThT-active amyloid fibrils (Fig. 5f).

## Discussion

In this work, we showed that domain-specific electrostatic interactions between PrP and α-Syn at a narrow stoichiometry regime result in the formation of highly dynamic liquid-like droplets with a mobile internal organization (Fig. 6). The charge neutralization drives the formation of these condensates, whereas, the charge inversion promotes their dispersion into a homogeneous solution akin to RNA-induced reentrant behavior. The entropic gain associated with counterion release upon LLPS allows these coacervates to display an LCST phase behavior. We demonstrated the critical role of different domains in promoting LLPS by using deletion mutations that revealed that the N-terminal disordered segment of PrP and the C-terminal domain of α-Syn are the principal drivers of two-component LLPS. Our site-specific picosecond time-resolved fluorescence anisotropy measurements revealed the formation of relatively ordered electrostatic nanoclusters that are stable on the nanosecond timescale. These clusters can act as oligomeric subunits connected via physical crosslinks within the condensed phase (Fig. 6). Our results also underscore the importance of timescales of breaking-and-making of non-covalent interactions in governing the hierarchical architecture and internal material property at different length scales. On the nanosecond timescale and molecular-to-nano length-scale, there can be a considerable structural and dynamical heterogeneity, whereas, on a much slower (seconds) timescale and mesoscopic length-scale, these assemblies display typical liquid-like characteristics. Such characteristics might be generic for a wide variety of multi-component condensates.

**Fig. 6.**
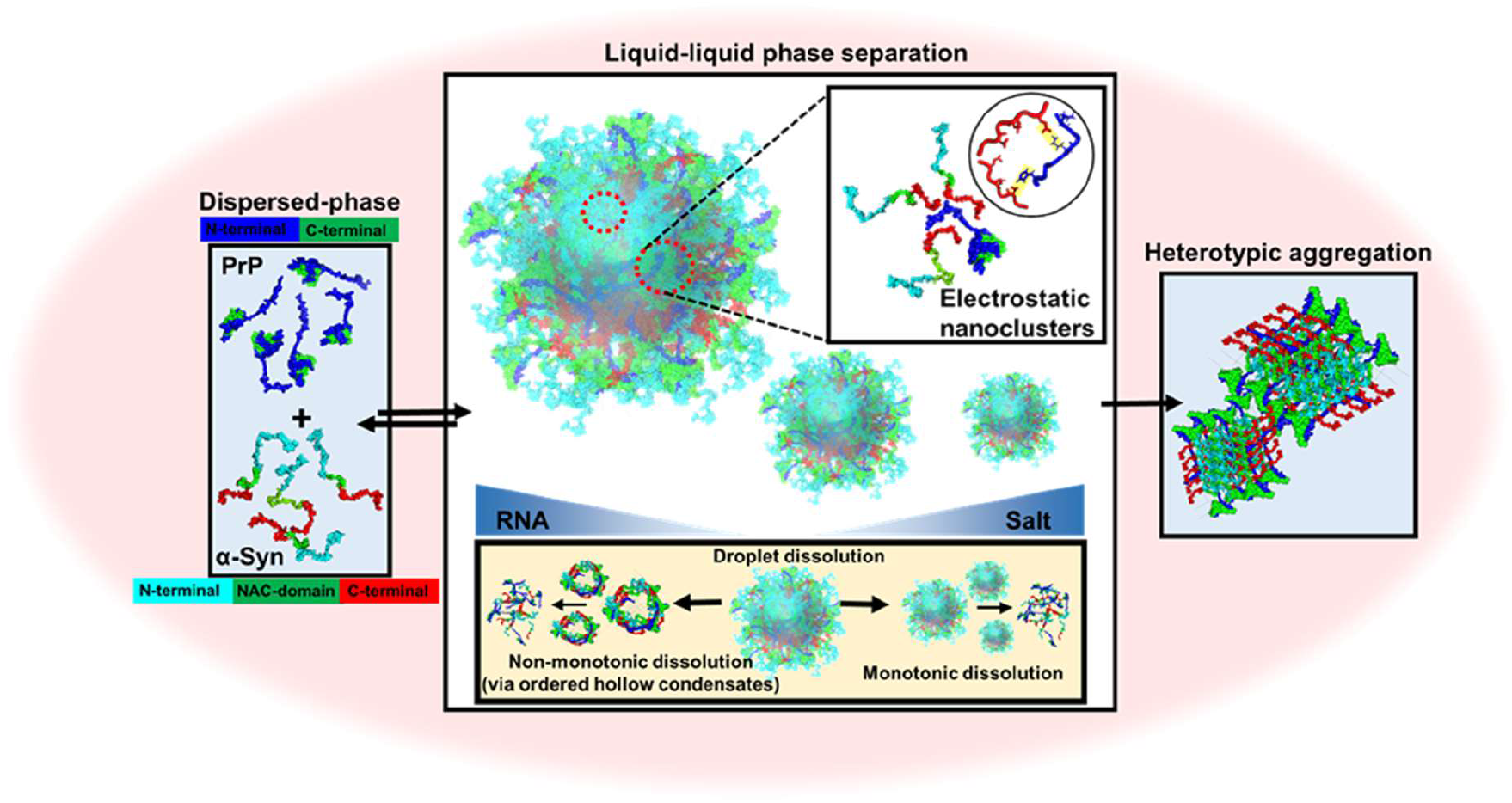
A schematic of PrP-α-Syn-RNA multicomponent condensates. Complex coacervation of PrP and α-Syn drives the formation of partially ordered electrostatic nanoclusters. The addition of salt results in a monotonic condensate dissolution, whereas, the addition of RNA results in a non-monotonic dissolution via multiphasic hollow condensates. PrP-α-Syn condensates undergo a liquid-to-solid transition into heterotypic amyloids.

In summary, our results demonstrate the crucial role of phase separation in modulating the interactions between PrP and α-Syn. Both PrP and α-Syn have been shown to be localized in lipid rafts.^78,79^ Although PrP is a well-folded GPI-anchored cell surface protein, its intrinsically disordered N-terminal segment retains a high degree of flexibility and multivalency that enables promiscuous interactions with multiple binding partners. IDR-enabled LLPS is emerging as one of the potential mechanisms for assembling cell-surface receptors to facilitate signal transduction. ^80–82^ Given the presence of both PrP and α-Syn within the lipid rafts, we speculate that the receptor clustering in lipid rafts might be one of the potential mechanisms to promote complex coacervation of PrP and α-Syn in the cellular context. Our findings also demonstrate the buffering role of RNA in their interactions. The addition of RNA to these preformed condensates weakens the α-Syn-PrP interaction and disrupts the formation of the ordered domains. Lower RNA concentrations yielded ternary droplets with an uniform distributions, whereas higher RNA concentrations resulted in the formation of ordered, multiphasic, vesicle-like condensates with charge-driven hierarchal architecture (Fig. 6). Such a concept of heterotypic buffering might potentially act as a framework to understand competitive protein-protein and protein-RNA interactions within the cellular milieu.^69,83^

The interaction of α-Syn with the N-terminal IDR of PrP is reminiscent of the recruitment of amyloid-β oligomers by PrP involved in synaptic impairment hinting at a plausible role of heterotypic α-Syn-PrP interactions in neurotoxicity.^84^ Recent studies have also provided evidence for the role of PrP as a membrane-surface receptor in the context of α-Syn oligomers and fibrils.^43–45^ The interaction elicits neurotoxic signaling pathways through metabotropic glutamate receptor 5, Fyn kinase, and N-methyl D-aspartate receptors resulting in synaptic dysfunction. Additionally, the presence of extensive α-Syn deposits has been linked to unique CJD cases with longer disease courses.^44,85^ Colocalization of α-Syn and PrP aggregates has been observed in early cytoplasmic inclusions.^42^ Our findings showcase the synergistic effect of PrP and α-Syn interactions in complex biomolecular condensates that can act as reaction crucibles to catalyze aberrant liquid-to-solid phase transitions into early heterotypic amyloids. Our study also highlights the pertinent role of LLPS in driving interactions between PrP and α-Syn, which under normal cellular conditions may be reversible; however, cellular stress can promote their aberrant phase transitions. Interestingly, such heterotypic interactions might potentially be involved in blocking prion propagation as shown previously.^44^ Furthermore, the heterotypic buffering in the presence of RNA can offer an alternate pathway in regulating their phase behavior and might be important in suppressing their potentially toxic effects. It is important to note that both PrP and α-Syn are transported via extracellular vesicles and exosomes which are highly enriched in RNA and perhaps can act as potential sites for such multiphasic interactions.^86–88^ The interplay of these critical molecular events can have broad implications in physiologically relevant receptor-mediated signaling as well as in disease-associated aberrant phase transitions.

## Methods

### Bioinformatic analyses of prion protein and α-Synuclein using various prediction tools

Classification of Intrinsically Disordered Ensemble Regions (CIDER)^53^ (http://pappulab.wustl.edu/CIDER/analysis) was used to predict the charge distribution and net charge per residue (NCPR) of human PrP and α-Syn. IUPred2a (https://iupred2a.elte.hu/) ^89^ was used to predict the intrinsic disorder. FuzPred/FuzDrop (http://protdyn-fuzpred.org/)^54^ and catGRANULE (http://s.tartaglialab.com)^55^ were used to predict the LLPS propensity of both PrP and α-Syn. All these data were plotted using Origin.

### Site-directed mutagenesis, protein expression, and purification

Recombinant full-length human PrP (PrP 23-231) plasmid cloned in vector pRSET-B was transformed in BL21(DE3)pLysS. PrP 23-144 (Y145Stop), PrP 112-231, and single cysteine variants of full-length PrP (W31C and W99C) were used as described previously.^63,90^ Recombinant full-length human α-Syn (1-140) plasmid cloned in vector pT7.7 was transformed in BL21(DE3)pLysS.^91^ α-Syn (1-102) (N103Stop) and α-Syn (1-132) (Y133Stop) were created using the full-length α-Syn plasmid. The primers used for α-Syn mutations are listed in Table S1. Single cysteine variants of α-Syn (A18C, A90C, and A124C) were used as described previously.^91^ Recombinant prion protein constructs were overexpressed and purified using previously published protocols.^63,90^ The purified proteins were refolded using the PD10 column in 14 mM MES buffer, pH 6.8. Recombinant α-Syn constructs except 103Stop were purified using the anion-exchange chromatography as described previously.^91^ α-Syn 103Stop was purified using cation-exchange chromatography. The purified proteins were dialyzed overnight for buffer exchange (14 mM MES buffer, pH 6.8). The purity of all the proteins was confirmed by SDS-PAGE analysis. Protein concentrations were estimated using ε_280 nm_ = 56,590 M^-1^cm^-1^ for PrP(23-231), ε_280 nm_ = 43,670 M^-1^cm^-1^ for Y145Stop, ε_280 nm_ = 14,200 M^-1^cm^-1^ for PrP (112-231), ε_278 nm_ = 5600 M^-1^cm^-1^ for wt α-Syn, ε_278 nm_ = 3840 M^-1^cm^-1^ for α-Syn 133Stop, ε_278 nm_ = 1280 M^-1^cm^-1^ for α-Syn 103Stop. All the experiments were performed using freshly purified proteins.

### Fluorescence labeling

Labeling of single cysteine variants were performed in denaturation buffer at pH 7.5 using thiol-active fluorescent dyes namely, fluorescein-5-maleimide (F5M), AlexaFluor488-C5-maleimide (Alexa-488), AlexaFluor594-C5-maleimide (Alexa-594), and 5-((2-((iodoacetyl)amino)ethyl)amino)naphthalene-1-sulfonic acid (IAEDANS). Excess free dye was removed using a PD10 desalting column. Labeled protein concentrations were estimated using molar extinction coefficients of the dyes [ε_495 nm_ = 68,000 M^-1^cm^-1^, for F5M; ε_493 nm_ = 72,000 M^-1^cm^-1^, for Alexa-488; ε_588 nm_ = 96,000 M^-1^cm^-1^ for Alexa-594 and ε_337 nm_ = 6100 M^-1^cm^-1^ for IAEDANS].

### Dynamic light scattering (DLS) and electrophoretic mobility measurements

Hydrodynamic radii were estimated using the DLS instrument (Malvern Zetasizer). All the buffers were filtered through 0.02 µm filters before measurements. Monomeric PrP (50 µM), α-Syn (50 µM) and droplets (PrP + α-Syn; 20 µM + 30 µM) were used for the measurements. Electrophoretic mobility measurements were estimated at 25 °C using the DLS instrument using the M3-PALS (Phase Analysis Light Scattering) method.

### Sedimentation assays

For sedimentation assays, complex coacervates (100 µL) of PrP (20 µM) and α-Syn (30 µM) were formed and incubated for 5 minutes. They were then centrifuged at 25,000 x g, 25 °C to separate dense phase and light phase. The supernatant was carefully removed, and the pellet (dense phase) obtained after centrifugation was resuspended in 8M urea (10 µL). The samples were run on an SDS-PAGE (15%) and were visualized using the Coomassie blue staining. The saturation concentration (C_sat_) was estimated by comparing the supernatant intensity with a known concentration intensity using ImageJ software. Hollow condensates of α-Syn-PrP in the presence of RNA (150 ng/µL) were also processed similarly, centrifuged, and analyzed using SDS-PAGE.

### Confocal microscopy

All the imaging experiments were performed at room temperature on ZEISS LSM 980 Elyra 7 super-resolution microscope equipped with a high-resolution monochrome cooled AxioCamMRm Rev. 3 FireWire(D) camera, using a 63x oil-immersion objective (Numerical aperture 1.4). For visualizing droplets of PrP-α-Syn and PrP-α-Syn-RNA complexes, 1% of unlabeled proteins were doped with labeled proteins, and the samples were placed in Labtek chambers. Alexa-488-labeled protein was imaged using a 488-nm laser diode (11.9 mW), and Alexa-594-labeled protein was imaged using an excitation source at 590-nm. The ThT positive fibrils were imaged using a 402-nm excitation source. Images were processed and analyzed using ImageJ (NIH, Bethesda, USA). Concentrations in the dense phase and the light phase were estimated using a previously defined protocol.^75^ Calibration plots were generated from fluorescence intensities of the dispersed phase of Alexa-488-labeled α-Syn and Alexa-594-labeled PrP at different concentrations. The confocal images of droplets formed using Alexa-488-labeled PrP, and Alexa-594-labeled α-Syn (labeled protein: 0.1%) were then analyzed using ImageJ software to get an approximate estimation of protein concentration inside droplets. The C_sat_ was estimated using a similar method (labeled protein: 2%) and was verified using SDS-PAGE analysis as described above.

### Fluorescence Recovery after Photobleaching (FRAP)

FRAP experiments were performed on ZEISS LSM 980 Elyra 7 super-resolution microscope equipped with a high-resolution monochrome cooled AxioCamMRm Rev. 3 FireWire(D) camera, using a 63x oil-immersion objective (Numerical aperture 1.4). Alexa-488-labeled α-Syn and PrP (∼ 1 %) were used for FRAP experiments. A region of interest (ROI) with a radius of 0.5 μm was bleached using a 488-nm laser. The recovery of the bleached spots was recorded using ZEN Pro 2011(ZEISS) software provided with the instrument. The fluorescence recovery curves were background corrected, normalized, and plotted using Origin.

### Phase separation assays

Phase separation was induced by mixing α-Syn, PrP, and RNA in the desired stoichiometries. The turbidity of the phase-separated samples for PrP-α-Syn-RNA complex (25 °C), PrP-RNA coacervates (25 °C), and PrP-wild-type with α-Syn/103Stop/133Stop (37 °C) were measured at 350 nm on Genova Life Science spectrophotometer (ver.1.51.4). The mean and the standard error were obtained from at least three independent sets of experiments for all the measurements. The turbidity of the phase-separated samples for α-Syn-wild-type with PrP (23-144)/ PrP (112-231) (37 °C) and PrP-α-Syn at different temperatures were estimated by taking absorbance at 350 nm on a Multiskan Go (Thermo scientific) plate reader using 96-well NUNC optical bottom plates. For temperature-dependent turbidity assays, the LLPS-induced solution was incubated for 5 min at respective temperatures to minimize any discrepancy due to temperature fluctuation. The sample volume used for these measurements was 150 μL, and raw turbidity data are plotted with background subtraction. For most of the experiments, the PrP concentration was fixed to 20 μM, and the α-Syn concentration was fixed at 30 μM, pH 6.8 unless otherwise mentioned.

### Circular dichroism spectroscopy (CD)

The far-UV CD experiments for monomeric proteins and droplets were recorded on a Chirascan CD spectrometer (Applied Photophysics, UK) at room temperature in a 1mm pathlength quartz cuvette. The data were recorded using final protein concentration as follows: PrP monomer: 20 µM, α-Syn monomer: 30 µM and droplets, PrP + α-Syn (20 µM + 30 µM). The aggregates of PrP-α-Syn (20 µM + 30 µM) and α-Syn were centrifuged at 25,000 x g, 37 °C to separate the monomeric population. The CD spectra were acquired after resuspending the pellet in 20 mM phosphate buffer, pH 7.5 on a Biologic MOS500 spectrometer. All the spectra were averaged over three scans and were blank subtracted which were then processed and plotted using Origin 2018.

### Aggregation kinetics

The thioflavin T (ThT) aggregation kinetics were performed using NUNC 96-well plate on POLARstar Omega Plate Reader Spectrophotometer (BMG LABTECH, Germany) at 37 °C. The reaction mixtures (150 µL) containing a glass bead were subjected to stirring conditions, 600 rpm, with protein concentrations mentioned in the respective plots. The final concentration of ThT in the reaction mixture was 20 μM. For bulk Raman measurements, similar aggregation reactions were set up without ThT in the reaction mixture. The kinetic traces were plotted using Origin.

### Steady-state fluorescence anisotropy

The steady-state fluorescence experiments were performed on a FluoroMax-4 spectrofluorometer (Horiba Jobin Yvon, NJ, USA) using a 1-mm pathlength quartz cuvette. Fluorescence measurements were performed for α-Syn-PrP condensates at 37 °C and α-Syn-PrP-RNA condensates at 25 °C. The concentrations of PrP and α-Syn were fixed at 20 µM and 30 µM, respectively. For recording fluorescein fluorescence, F5M-labeled PrP and α-Syn (200 nM of labeled protein mixed with the unlabeled protein) were used. The samples were excited at 485 nm and the emission spectra were collected in the range between 510 nm and 600 nm. For recording fluorescence of IAEDANS-labeled single-Cys 124 of α-Syn (15 µM of labeled protein was mixed with the wild-type protein), the samples were excited at 375 nm, and the emission spectra were collected in the range between 400 nm and 600 nm. The steady-state fluorescence anisotropy was recorded at emission maxima. The steady-state fluorescence anisotropy (*r*_ss_) is the estimated form of the following relationship.

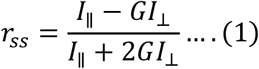

where *I*_∥_ and *I*_⊥_ are the parallel and perpendicular fluorescence intensities, respectively, and the measured intensities were corrected using the G-factor.

### Picosecond time-resolved fluorescence anisotropy measurements

Time-resolved fluorescence anisotropy decay measurements were performed using a time-correlated single-photon counting (TCSPC) set up (Horiba Jobin Yvon, NJ). The samples were excited using 485-nm and 375-nm NanoLED picosecond laser diodes for F5M and IAEDANS labeled proteins, respectively. The instrument response function (IRF) was obtained by using a dilute solution of colloidal silica (Ludox) and the full-width half maxima (FWHM) was estimated to be ∼265 ps. All the measurements were performed at 37 °C. To record anisotropy decay profiles, the emission wavelength was set at the respective emission maxima, with a bandpass of 8 nm. The fluorescence intensities were collected at 0° (*I*_║_) and 90° (*I*_*┴*_) with respect to the geometric orientation of the excitation polarizer. The perpendicular fluorescence intensity decays were corrected using the *G*-factor obtained from free dyes in water. The anisotropy decays were analyzed by global fitting of *I*_*┴*_ and *I*_║_ using the following equation:

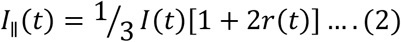

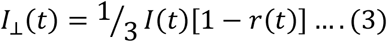

where *I* represent the time-dependent fluorescence intensity collected at the magic angle (54.7°). The time-resolved depolarization kinetics is described using a typical biexponential decay function which defines fast (*ϕ*_1_) and slow (*ϕ*_2_) rotational correlation times arising due to local motion of the fluorophore and segmental mobility of IDP, respectively.^66^

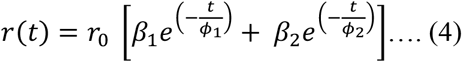

where *r*_0_ represents the intrinsic time zero or the fundamental anisotropy of the attached fluorophore. *β*_1_ and *β*_2_ represent the amplitudes associated with fast and slow rotational correlation time, respectively. The goodness of fit was evaluated by reduced *χ*^2^ values, the randomness of the residuals, and autocorrelation functions.^66^

For dispersed monomers and droplets (α-Syn residue 18), the anisotropy decays were fitted using a biexponential equation. However, for droplets (α-Syn residue 90, 124; PrP residue 31 and 99), time-resolved anisotropy decays could only be described by a triexponential decay kinetics with an additional slower correlation time (*ϕ*_3_) as shown below.

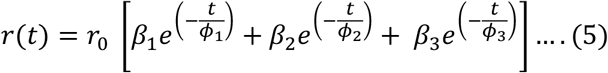

In order to get a better estimate of the slow correlation time (*ϕ*_3_) corresponding to the electrostatic clusters, a longer lifetime label (IAEDANS with a mean lifetime of 12 ns) was used for the measurements. The *ϕ*_3_ was found to be 54 ± 6 ns (For recovered parameters, see Table S2). This value was used for estimating the approximate hydrodynamic radii (*R*_h_) of the nanoclusters. The *R*_h_ was estimated using the Stokes-Einstein equation as follows.

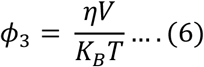

where *η* is the viscosity of the medium, *V* is the volume of the rotating unit 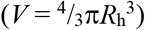, *k*_*B*_ is the Boltzmann constant, and *T* is the temperature. The robustness of the recovered correlation time (*ϕ*_3_) was also assessed by using both free and forced fits.

### Raman spectroscopy

Raman spectra for heterotypic PrP-α-Syn aggregates and homotypic α-Syn aggregates were recorded on an inVia laser Raman microscope (Renishaw, UK). The aggregates were centrifuged at 25,000 x g, and the obtained pellets were resuspended in 5 µL of 20 mM phosphate buffer, pH 7.5. The sample volume of 5 µL was deposited and dried onto a glass slide covered with an aluminum sheet. The sample was focused using a 100x objective lens (Nikon, Japan), and a 785-nm 500 mW NIR laser with a 50% laser power and exposure time of 10 s was used for excitation. The Rayleigh scattering was filtered by using an edge filter of 785-nm. The Raman scattering was collected and dispersed using a 1200 lines/mm diffraction grating and detected using an air-cooled CCD detector. Inbuilt Wire 3.4 software was used for data acquisition. All the data were averaged over 20 scans. Baseline correction and smoothening of the acquired spectra were performed using Wire 3.4 and the spectra were plotted using Origin.

### Atomic Force Microscopy (AFM)

AFM images of PrP-α-Syn co-aggregates were acquired on Innova atomic force microscope (Bruker) operating in tapping mode. For sample preparation, 10 μL of the aliquots were taken from the reaction mixture and were deposited onto the freshly cleaved, Milli-Q water-washed muscovite mica (Grade V-4 mica from SPI, PA). The samples were incubated for 5 minutes at room temperature and were washed twice with 100 μL of filtered Milli-Q water. The samples were further dried under a gentle stream of nitrogen before AFM imaging. NanoDrive (v8.03) software was used for the data acquisition and the acquired images were processed using WSxM 5.0D 8.1 software.^92^ The height profiles were obtained from WSxM software and were plotted using Origin.

## Acknowledgments

We thank IISER Mohali, Department of Science and Technology (Nano-Mission to S.M. and FIST grant to the Department of Biological Sciences, IISER Mohali), Department of Biotechnology (fellowship to A.A.), Council of Scientific and Industrial Research (fellowship to S.K.R.), Ministry of Education, Govt. of India (Centre of Excellence grant to S.M.) for financial support, Prof. Witold Surewicz (Case Western Reserve University, USA) and Prof. Vinod Subramaniam (University of Twente, Netherlands) for the kind gift of the DNA plasmids for full-length PrP and α-Syn, respectively, Prof. N. Periasamy (Retd. TIFR Mumbai) for providing us with the fluorescence decay analysis program, Ms. Debapriya Das for helping with fluorescence decay analysis, and Dr. Mily Bhattacharya (Thapar Institute) and the members of the Mukhopadhyay lab for critically reading this manuscript.

## References

1. Alberti, S. and Hyman, A.A., 2021. Biomolecular condensates at the nexus of cellular stress, protein aggregation disease and ageing. Nature Reviews Molecular Cell Biology, 22(3), pp.196–213.

2. Lyon, A.S., Peeples, W.B. and Rosen, M.K., 2021. A framework for understanding the functions of biomolecular condensates across scales. Nature Reviews Molecular Cell Biology, 22(3), pp.215–235.

3. Fuxreiter, M. and Vendruscolo, M., 2021. Generic nature of the condensed states of proteins. Nature Cell Biology, 23(6), pp.587–594.

4. Roden, C. and Gladfelter, A.S., 2021. RNA contributions to the form and function of biomolecular condensates. Nature Reviews Molecular Cell Biology, 22(3), pp.183–195.

5. Choi, J.M., Holehouse, A.S. and Pappu, R.V., 2020. Physical principles underlying the complex biology of intracellular phase transitions. Annual Review of Biophysics, 49, pp.107–133.

6. Boeynaems, S., Alberti, S., Fawzi, N.L., Mittag, T., Polymenidou, M., Rousseau, F., Schymkowitz, J., Shorter, J., Wolozin, B., Van Den Bosch, L. and Tompa, P., 2018. Protein phase separation: a new phase in cell biology. Trends in cell biology, 28(6), pp.420–435.

7. Mitrea, D.M. and Kriwacki, R.W., 2016. Phase separation in biology; functional organization of a higher order. Cell Communication and Signaling, 14(1), pp.1–20.

8. Hyman, A.A., Weber, C.A. and Jülicher, F., 2014. Liquid-liquid phase separation in biology. Annual review of cell and developmental biology, 30, pp.39–58.

9. Garabedian, M.V., Wang, W., Dabdoub, J.B., Tong, M., Caldwell, R.M., Benman, W., Schuster, B.S., Deiters, A. and Good, M.C., 2021. Designer membraneless organelles sequester native factors for control of cell behavior. Nature Chemical Biology, pp.1–10.

10. Fare, C.M., Villani, A., Drake, L.E. and Shorter, J., 2021. Higher-order organization of biomolecular condensates. Open biology, 11(6), p.210137.

11. Banerjee, P.R., Milin, A.N., Moosa, M.M., Onuchic, P.L. and Deniz, A.A., 2017. Reentrant phase transition drives dynamic substructure formation in ribonucleoprotein droplets. Angewandte Chemie, 129(38), pp.11512–11517.

12. Korkmazhan, E., Tompa, P. and Dunn, A.R., 2021. The role of ordered cooperative assembly in biomolecular condensates. Nature Reviews Molecular Cell Biology, 1–2.

13. Lafontaine, D.L., Riback, J.A., Bascetin, R. and Brangwynne, C.P., 2021. The nucleolus as a multiphase liquid condensate. Nature Reviews Molecular Cell Biology, 22(3), pp.165–182.

14. Banani, S.F., Rice, A.M., Peeples, W.B., Lin, Y., Jain, S., Parker, R. and Rosen, M.K., 2016. Compositional control of phase-separated cellular bodies. Cell, 166(3), pp.651–663.

15. Ditlev, J.A., Case, L.B. and Rosen, M.K., 2018. Who’s in and who’s out—compositional control of biomolecular condensates. Journal of molecular biology, 430(23), pp.4666–4684.

16. Dignon, G.L., Zheng, W., Kim, Y.C., Best, R.B. and Mittal, J., 2018. Sequence determinants of protein phase behavior from a coarse-grained model. PLoS computational biology, 14(1), p.e1005941.

17. Vernon, R.M., Chong, P.A., Tsang, B., Kim, T.H., Bah, A., Farber, P., Lin, H. and Forman-Kay, J.D., 2018. Pi-Pi contacts are an overlooked protein feature relevant to phase separation. elife, 7, p.e31486.

18. Murthy, A.C., Dignon, G.L., Kan, Y., Zerze, G.H., Parekh, S.H., Mittal, J. and Fawzi, N.L., 2019. Molecular interactions underlying liquid− liquid phase separation of the FUS low-complexity domain. Nature structural & molecular biology, 26(7), pp.637–648.

19. Pak, C.W., Kosno, M., Holehouse, A.S., Padrick, S.B., Mittal, A., Ali, R., Yunus, A.A., Liu, D.R., Pappu, R.V. and Rosen, M.K., 2016. Sequence determinants of intracellular phase separation by complex coacervation of a disordered protein. Molecular Cell, 63(1), pp.72–85.

20. Riback, J.A., Zhu, L., Ferrolino, M.C., Tolbert, M., Mitrea, D.M., Sanders, D.W., Wei, M.T., Kriwacki, R.W. and Brangwynne, C.P., 2020. Composition-dependent thermodynamics of intracellular phase separation. Nature, 581(7807), pp.209–214.

21. Uversky, V.N., 2017. Intrinsically disordered proteins in overcrowded milieu: Membrane-less organelles, phase separation, and intrinsic disorder. Current opinion in structural biology, 44, pp.18–30.

22. Dignon, G.L., Best, R.B. and Mittal, J., 2020. Biomolecular phase separation: From molecular driving forces to macroscopic properties. Annual review of physical chemistry, 71, pp.53–75.

23. Martin, E.W. and Mittag, T., 2018. Relationship of sequence and phase separation in protein low-complexity regions. Biochemistry, 57(17), pp.2478–2487.

24. Martin, E.W., Holehouse, A.S., Peran, I., Farag, M., Incicco, J.J., Bremer, A., Grace, C.R., Soranno, A., Pappu, R.V. and Mittag, T., 2020. Valence and patterning of aromatic residues determine the phase behavior of prion-like domains. Science, 367(6478), pp.694–699.

25. Ruff, K.M., Roberts, S., Chilkoti, A. and Pappu, R.V., 2018. Advances in understanding stimulus-responsive phase behavior of intrinsically disordered protein polymers. Journal of molecular biology, 430(23), pp.4619–4635.

26. Brangwynne, C.P., Tompa, P. and Pappu, R.V., 2015. Polymer physics of intracellular phase transitions. Nature Physics, 11(11), pp.899–904.

27. Quiroz, F.G. and Chilkoti, A., 2015. Sequence heuristics to encode phase behaviour in intrinsically disordered protein polymers. Nature materials, 14(11), pp.1164–1171.

28. Saar, K.L., Morgunov, A.S., Qi, R., Arter, W.E., Krainer, G. and Knowles, T.P., 2021. Learning the molecular grammar of protein condensates from sequence determinants and embeddings. Proceedings of the National Academy of Sciences, 118(15).

29. Ranganathan, S. and Shakhnovich, E.I., 2020. Dynamic metastable long-living droplets formed by sticker-spacer proteins. Elife, 9, p.e56159.

30. Martin, E.W. and Mittag, T., 2018. Relationship of sequence and phase separation in protein low-complexity regions. Biochemistry, 57(17), pp.2478–2487.

31. Siegert, A., Rankovic, M., Favretto, F., Ukmar-Godec, T., Strohäker, T., Becker, S. and Zweckstetter, M., 2021. Interplay between tau and α-synuclein liquid–liquid phase separation. Protein Science, 30(7), pp.1326–1336.

32. Bhopatkar, A.A., Uversky, V.N. and Rangachari, V., 2020. Granulins modulate liquid– liquid phase separation and aggregation of the prion-like C-terminal domain of the neurodegeneration-associated protein TDP-43. Journal of Biological Chemistry, 295(8), pp.2506–2519.

33. Ray, S., Singh, N., Kumar, R., Patel, K., Pandey, S., Datta, D., Mahato, J., Panigrahi, R., Navalkar, A., Mehra, S. and Gadhe, L.,… Maji, S. K. 2020. α-Synuclein aggregation nucleates through liquid–liquid phase separation. Nature Chemistry, 12(8), pp.705–716.

34. Elbaum-Garfinkle, S. and Kriwacki, R.W., 2021. Phase Separation in Biology & Disease: The Next Chapter. Journal of Molecular Biology, pp.166990–166990.

35. Aguzzi, A. and Altmeyer, M., 2016. Phase separation: linking cellular compartmentalization to disease. Trends in cell biology, 26(7), pp.547–558.

36. Alberti, S. and Dormann, D., 2019. Liquid–liquid phase separation in disease. Annual review of genetics, 53, pp.171–194.

37. Katorcha, E., Makarava, N., Lee, Y.J., Lindberg, I., Monteiro, M.J., Kovacs, G.G. and Baskakov, I.V., 2017. Cross-seeding of prions by aggregated α-synuclein leads to transmissible spongiform encephalopathy. PLoS pathogens, 13(8), p.e1006563.

38. Kovacs, G.G., Rahimi, J., Ströbel, T., Lutz, M.I., Regelsberger, G., Streichenberger, N., Perret-Liaudet, A., Höftberger, R., Liberski, P.P., Budka, H. and Sikorska, B., 2017. Tau pathology in Creutzfeldt-Jakob disease revisited. Brain Pathology, 27(3), pp.332–344.

39. Irwin, D.J., Lee, V.M.Y. and Trojanowski, J.Q., 2013. Parkinson’s disease dementia: convergence of α-synuclein, tau and amyloid-β pathologies. Nature Reviews Neuroscience, 14(9), pp.626–636.

40. Goedert, M., 2015. Alzheimer’s and Parkinson’s diseases: The prion concept in relation to assembled Aβ, tau, and α-synuclein. Science, 349(6248).

41. Nakashima-Yasuda, H., Uryu, K., Robinson, J., Xie, S.X., Hurtig, H., Duda, J.E., Arnold, S.E., Siderowf, A., Grossman, M., Leverenz, J.B. and Woltjer, R., 2007. Co-morbidity of TDP-43 proteinopathy in Lewy body related diseases. Acta neuropathologica, 114(3), pp.221–229.

42. Kovacs, G.G., Zerbi, P., Voigtländer, T., Strohschneider, M., Trabattoni, G., Hainfellner, J.A. and Budka, H., 2002. The prion protein in human neurodegenerative disorders. Neuroscience letters, 329(3), pp.269–272.

43. Brás, I.C., Lopes, L.V. and Outeiro, T.F., 2018. Sensing α-Synuclein From the Outside via the Prion Protein: Implications for Neurodegeneration. Movement Disorders, 33(11), pp.1675–1684.

44. Aulić, S., Masperone, L., Narkiewicz, J., Isopi, E., Bistaffa, E., Ambrosetti, E., Pastore, B., De Cecco, E., Scaini, D., Zago, P. and Moda, F., 2017. α-Synuclein amyloids hijack prion protein to gain cell entry, facilitate cell-to-cell spreading and block prion replication. Scientific reports, 7(1), pp.1–12.

45. Rösener, N.S., Gremer, L., Wördehoff, M.M., Kupreichyk, T., Etzkorn, M., Neudecker, P. and Hoyer, W., 2020. Clustering of human prion protein and α-synuclein oligomers requires the prion protein N-terminus. Communications biology, 3(1), pp.1–12.

46. Ferreira, D.G., Temido-Ferreira, M., Miranda, H.V., Batalha, V.L., Coelho, J.E., Szegö, É.M., Marques-Morgado, I., Vaz, S.H., Rhee, J.S., Schmitz, M. and Zerr, I., 2017. α-synuclein interacts with PrP C to induce cognitive impairment through mGluR5 and NMDAR2B. Nature neuroscience, 20(11), pp.1569–1579.

47. Masliah, E., Rockenstein, E., Inglis, C., Adame, A., Bett, C., Lucero, M. and Sigurdson, C.J., 2012. Prion infection promotes extensive accumulation of α-synuclein in aged human α-synuclein transgenic mice. Prion, 6(2), pp.184–190.

48. Lashuel, H.A., Overk, C.R., Oueslati, A. and Masliah, E., 2013. The many faces of α-synuclein: from structure and toxicity to therapeutic target. Nature Reviews Neuroscience, 14(1), pp.38–48.

49. Prusiner, S.B., 1982. Novel proteinaceous infectious particles cause scrapie. Science, 216(4542), pp.136–144.

50. Prusiner, S.B., 2017. Prion biology. Cold Spring Harbour Laboratory Press.

51. Scheckel, C. and Aguzzi, A., 2018. Prions, prionoids and protein misfolding disorders. Nature Reviews Genetics, 19(7), pp.405–418.

52. Zahn, R., Liu, A., Lührs, T., Riek, R., von Schroetter, C., García, F.L., Billeter, M., Calzolai, L., Wider, G. and Wüthrich, K., 2000. NMR solution structure of the human prion protein. Proceedings of the National Academy of Sciences, 97(1), pp.145–150.

53. Holehouse, A.S., Ahad, J., Das, R.K. and Pappu, R.V., 2015. CIDER: Classification of intrinsically disordered ensemble regions. Biophysical Journal, 108(2), p.228a.

54. Hardenberg, M., Horvath, A., Ambrus, V., Fuxreiter, M. and Vendruscolo, M., 2020. Widespread occurrence of the droplet state of proteins in the human proteome. Proceedings of the National Academy of Sciences, 117(52), pp.33254–33262.

55. Bolognesi, B., Gotor, N.L., Dhar, R., Cirillo, D., Baldrighi, M., Tartaglia, G.G. and Lehner, B., 2016. A concentration-dependent liquid phase separation can cause toxicity upon increased protein expression. Cell reports, 16(1), pp.222–231.

56. Srivastava, S. and Baskakov, I.V., 2015. Contrasting effects of two lipid cofactors of prion replication on the conformation of the prion protein. PLoS One, 10(6), p.e0130283.

57. Jain, N., Bhasne, K., Hemaswasthi, M. and Mukhopadhyay, S., 2013. Structural and dynamical insights into the membrane-bound α-synuclein. PloS one, 8(12), p.e83752.

58. Chang, L.W., Lytle, T.K., Radhakrishna, M., Madinya, J.J., Vélez, J., Sing, C.E. and Perry, S.L., 2017. Sequence and entropy-based control of complex coacervates. Nature communications, 8(1), pp.1–8.

59. Pak, C.W., Kosno, M., Holehouse, A.S., Padrick, S.B., Mittal, A., Ali, R., Yunus, A.A., Liu, D.R., Pappu, R.V. and Rosen, M.K., 2016. Sequence determinants of intracellular phase separation by complex coacervation of a disordered protein. Molecular cell, 63(1), pp.72–85.

60. Park, S., Barnes, R., Lin, Y., Jeon, B.J., Najafi, S., Delaney, K.T., Fredrickson, G.H., Shea, J.E., Hwang, D.S. and Han, S., 2020. Dehydration entropy drives liquid-liquid phase separation by molecular crowding. Communications Chemistry, 3(1), pp.1–12.

61. Sorrentino, Z.A. and Giasson, B.I., 2020. The emerging role of α-synuclein truncation in aggregation and disease. Journal of Biological Chemistry, 295(30), pp.10224–10244.

62. Choi, J.K., Cali, I., Surewicz, K., Kong, Q., Gambetti, P. and Surewicz, W.K., 2016. Amyloid fibrils from the N-terminal prion protein fragment are infectious. Proceedings of the National Academy of Sciences, 113(48), pp.13851–13856.

63. Agarwal, A., Rai, S.K., Avni, A. and Mukhopadhyay, S., 2021. An Intrinsically Disordered Pathological Variant of the Prion Protein Y145Stop Transforms into Self-Templating Amyloids via Liquid-Liquid Phase Separation. bioRxiv (2021) 2021.01.09.426049.

64. Peixoto, P.D., Tavares, G.M., Croguennec, T., Nicolas, A., Hamon, P., Roiland, C. and Bouhallab, S., 2016. Structure and dynamics of heteroprotein coacervates. Langmuir, 32(31), pp.7821–7828.

65. Kizilay, E., Seeman, D., Yan, Y., Du, X., Dubin, P.L., Donato-Capel, L., Bovetto, L. and Schmitt, C., 2014. Structure of bovine β-lactoglobulin–lactoferrin coacervates. Soft Matter, 10(37), pp.7262–7268.

66. Majumdar, A. and Mukhopadhyay, S., 2018. Fluorescence depolarization kinetics to study the conformational preference, structural plasticity, binding, and assembly of intrinsically disordered proteins. Methods in enzymology, 611, pp.347–381.

67. Jain, N., Narang, D., Bhasne, K., Dalal, V., Arya, S., Bhattacharya, M. and Mukhopadhyay, S., 2016. Direct observation of the intrinsic backbone torsional mobility of disordered proteins. Biophysical journal, 111(4), pp.768–774.

68. Joseph R.. Lakowicz, 1999. Principles of fluorescence spectroscopy. Kluwer Academic/Plenum.

69. Sanders, D.W., Kedersha, N., Lee, D.S., Strom, A.R., Drake, V., Riback, J.A., Bracha, D., Eeftens, J.M., Iwanicki, A., Wang, A. and Wei, M.T., 2020. Competing protein-RNA interaction networks control multiphase intracellular organization. Cell, 181(2), pp.306–324.

70. Sanchez-Burgos, I., Espinosa, J.R., Joseph, J.A. and Collepardo-Guevara, R., 2021. Valency and binding affinity variations can regulate the multilayered organization of protein condensates with many components. Biomolecules, 11(2), p.278.

71. Gomes, M.P., Cordeiro, Y. and Silva, J.L., 2008. The peculiar interaction between mammalian prion protein and RNA. Prion, 2(2), pp.64–66.

72. Silva, J.L., Lima, L.M.T., Foguel, D. and Cordeiro, Y., 2008. Intriguing nucleic-acid-binding features of mammalian prion protein. Trends in biochemical sciences, 33(3), pp.132–140.

73. Alshareedah, I., Moosa, M.M., Raju, M., Potoyan, D.A. and Banerjee, P.R., 2020. Phase transition of RNA− protein complexes into ordered hollow condensates. Proceedings of the National Academy of Sciences, 117(27), pp.15650–15658.

74. Portz, B., Lee, B.L. and Shorter, J., 2021. FUS and TDP-43 phases in health and disease. Trends in biochemical sciences.

75. Wegmann, S., Eftekharzadeh, B., Tepper, K, Zoltowska, K. M., Bennett, R. E., Dujardin, S., Laskowski, P. R., MacKenzie, D., Kamath, T., Commins, C., Vanderburg, C., Roe, A. D., Fan, Z., Molliex, A. M., Hernandez-Vega, A., Muller, D., Hyman, A. A., Mandelkow, E., Taylor, J.P., and Hyman, B.T., 2018. Tau protein liquid-liquid phase separation can initiate tau aggregation. The EMBO journal, 37(7), p.e98049.

76. Haik, S., Privat, N., Adjou, K., Sazdovitch, V., Dormont, D., Duyckaerts, C. and Hauw, J., 2002. Alpha-synuclein-immunoreactive deposits in human and animal prion diseases. Acta neuropathologica, 103(5), pp.516–520.

77. Bhattacharya, M., Jain, N., Dogra, P., Samai, S. and Mukhopadhyay, S., 2013. Nanoscopic amyloid pores formed via stepwise protein assembly. The journal of physical chemistry letters, 4(3), pp.480–485.

78. Taylor, D.R. and Hooper, N.M., 2006. The prion protein and lipid rafts. Molecular membrane biology, 23(1), pp.89–99.

79. Fortin, D.L., Troyer, M.D., Nakamura, K., Kubo, S.I., Anthony, M.D. and Edwards, R.H., 2004. Lipid rafts mediate the synaptic localization of α-synuclein. Journal of Neuroscience, 24(30), pp.6715–6723.

80. Forman-Kay, J.D., Kriwacki, R.W. and Seydoux, G., 2018. Phase separation in biology and disease. Journal of molecular biology, 430(23), p.4603.

81. Chong, P.A. and Forman-Kay, J.D., 2016. Liquid–liquid phase separation in cellular signaling systems. Current opinion in structural biology, 41, pp.180–186.

82. Nesterov, S.V., Ilyinsky, N.S. and Uversky, V.N., 2021. Liquid-liquid phase separation as a common organizing principle of intracellular space and biomembranes providing dynamic adaptive responses. Biochimica et Biophysica Acta (BBA)-Molecular Cell Research, p.119102.

83. Kaur, T., Raju, M., Alshareedah, I., Davis, R.B., Potoyan, D.A. and Banerjee, P.R., 2021. Sequence-encoded and composition-dependent protein-RNA interactions control multiphasic condensate morphologies. Nature communications, 12(1), pp.1–16.

84. Brody, A.H. and Strittmatter, S.M., 2018. Synaptotoxic signaling by amyloid beta oligomers in Alzheimer’s disease through prion protein and mGluR5. Advances in Pharmacology, 82, pp.293–323.

85. D’Arcy, C.E., Bitnun, A., Coulthart, M.B., D’Amour, R., Friedman, J., Knox, J.D., Rapoport, A., Carter, S., Widjaja, E., Hazrati, L.N. and Jansen, G.H., 2019. Sporadic Creutzfeldt-Jakob disease in a young girl with unusually long survival. Journal of Neuropathology & Experimental Neurology, 78(4), pp.373–378.

86. Hartmann, A., Muth, C., Dabrowski, O., Krasemann, S. and Glatzel, M., 2017. Exosomes and the prion protein: more than one truth. Frontiers in neuroscience, 11, p.194.

87. Lemprière, S., 2020. Exosomal α-synuclein as a biomarker for Parkinson disease. Nature Reviews Neurology, 16(5), pp.242–243.

88. Zomer, A., Vendrig, T., Hopmans, E.S., van Eijndhoven, M., Middeldorp, J.M. and Pegtel, D.M., 2010. Exosomes: fit to deliver small RNA. Communicative & integrative biology, 3(5), pp.447–450.

89. Mészáros, B., Erdős, G. and Dosztányi, Z., 2018. IUPred2A: context-dependent prediction of protein disorder as a function of redox state and protein binding. Nucleic acids research, 46(W1), pp.W329–W337.

90. Madhu, P. and Mukhopadhyay, S., 2019. Preferential recruitment of conformationally distinct amyloid-β oligomers by the intrinsically disordered region of the human prion protein. ACS chemical neuroscience, 11(1), pp.86–98.

91. Majumdar, A., Das, D., Madhu, P., Avni, A. and Mukhopadhyay, S., 2020. Excitation Energy Migration Unveils Fuzzy Interfaces within the Amyloid Architecture. Biophysical journal, 118(11), pp.2621–2626.

92. Horcas, I., Fernández, R., Gomez-Rodriguez, J.M., Colchero, J.W.S.X., Gómez-Herrero, J.W.S.X.M. and Baro, A.M., 2007. WSXM: a software for scanning probe microscopy and a tool for nanotechnology. Review of scientific instruments, 78(1), p.013705.

