## Supporting Information for "Spatiotemporal Modulations in Heterotypic Condensates of Prion and α-Synuclein Control Phase Transitions and Amyloid Conversion"

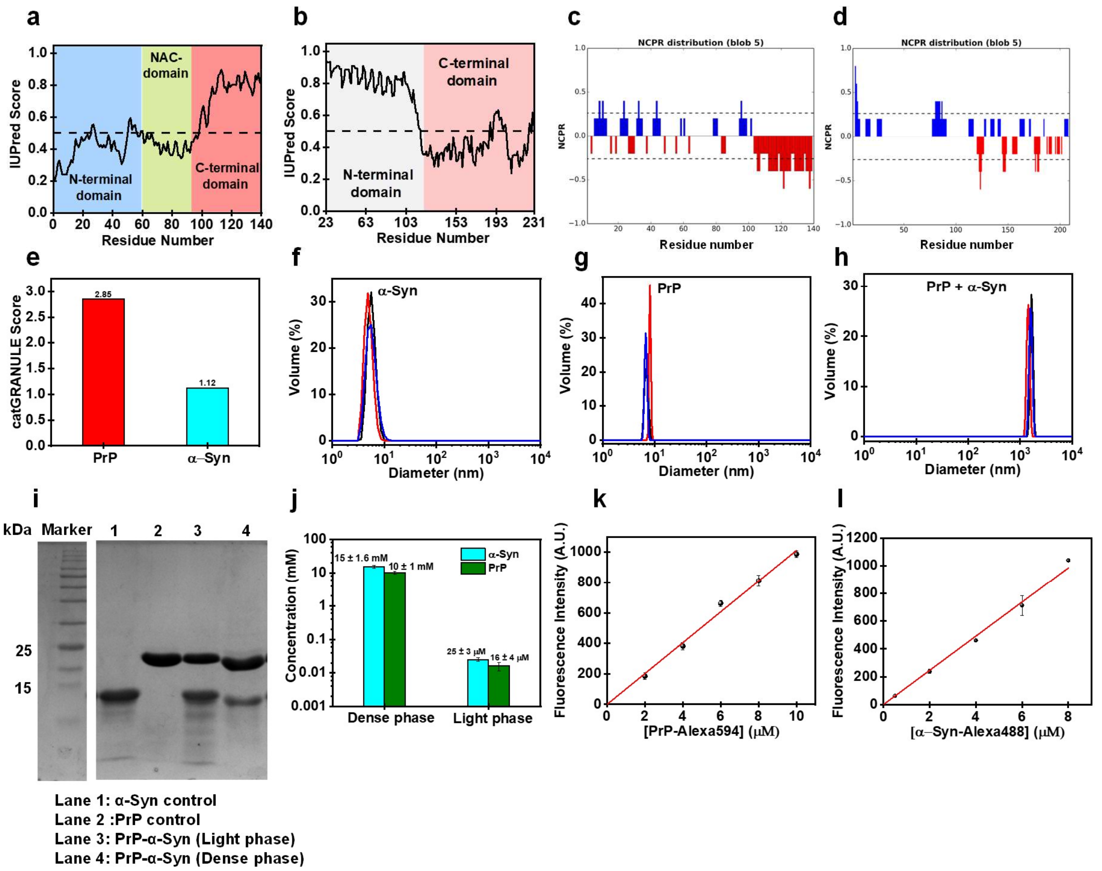


**Fig. S1** Prediction of intrinsic disorder using IUPred for **a** α-Syn and **b** full-length PrP (PrP 23-231). Net charge prediction tool CIDER indicating positively charged residues (blue) and negatively charged residues (red) for **c** α-Syn and **d** PrP (23-231). **e** Prediction of phase separation propensity of PrP and α-Syn using catGRANULE. Particle size distribution using dynamic light scattering for **f** α-Syn monomer (50 μM) **g** PrP monomer (50 μM) and **h** PrP-α-Syn droplets (20 μM + 30 μM). **i** SDS-PAGE for PrP-α-Syn droplets after sedimentation assay. **j-l** Dense phase and light phase concentrations estimation for PrP and α-Syn within droplets using fluorescence intensity calibration. The fluorescence intensity calibration plot using different concentrations of Alexa-488-labeled α-Syn and Alexa-594-labeled PrP.

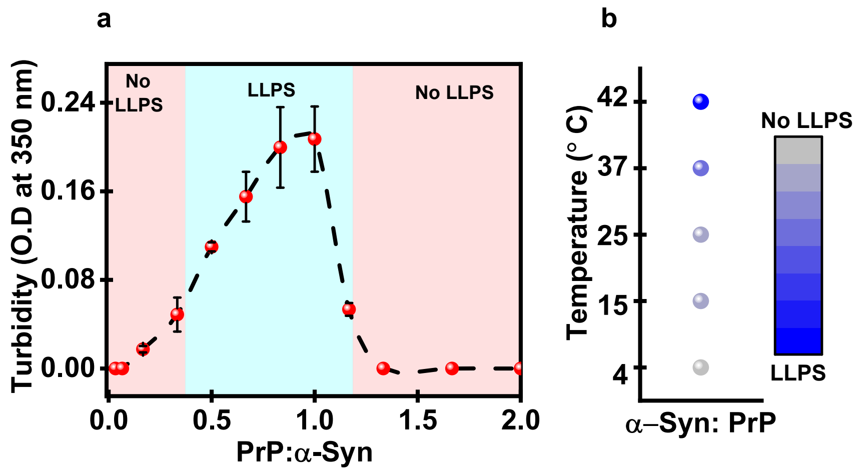


**Fig. S2 a** Solution turbidity plot at a fixed α-Syn concentration as a function of increasing PrP concentrations showing reentrant behavior. **b** Phase diagram for α-Syn:PrP (1.5) as a function of increasing temperature constructed from mean turbidity values.

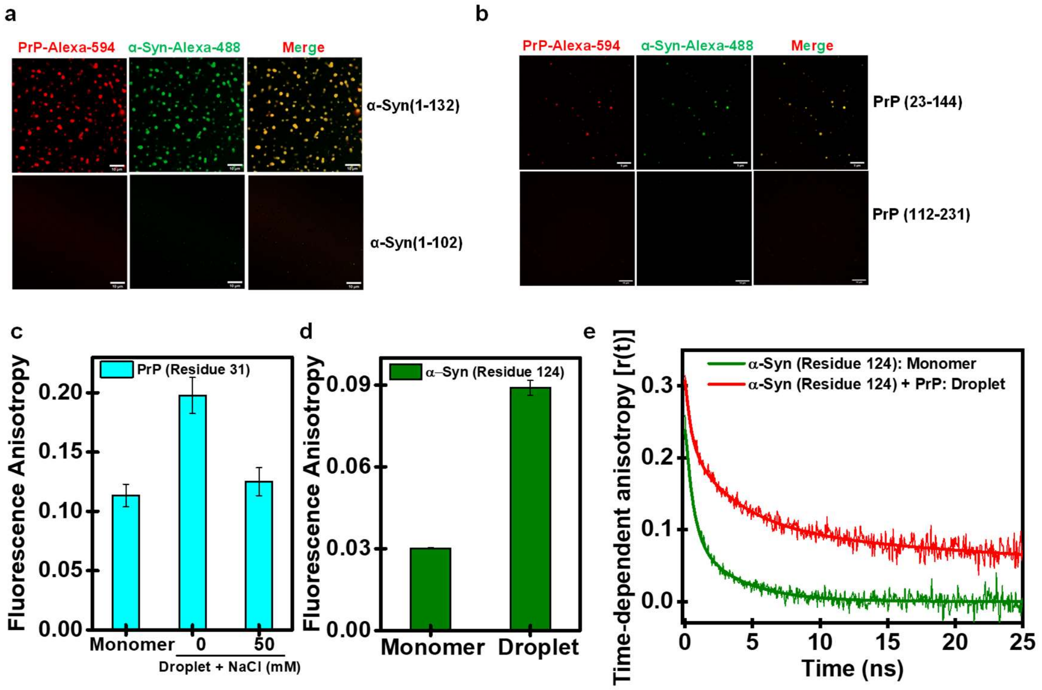


**Fig. S3 a** Confocal fluorescence images of Alexa-594-labeled PrP with different constructs of Alexa-488-labeled α-Syn. 20 μM of PrP and 60 μM α-Syn (1-132) and α-Syn (1-102) was used for imaging. Scale bar: 10 μm. **b** Confocal fluorescence images of Alexa-488-labeled α-Syn with different constructs of Alexa-594-labeled PrP. 20 μM of PrP and 50 μM of α-Syn were used for imaging. Scale bar: 10 μm. **c** Steady-state fluorescence anisotropy for F5M-labeled PrP residue 31 indicating droplet dissolution in the presence of NaCl. **d** Steady-state fluorescence anisotropy for IAEDANS-labeled α-Syn residue 124. **e** Time-resolved anisotropy decay of IAEDANS-labeled α-Syn residue 124 in dispersed monomer and droplets.

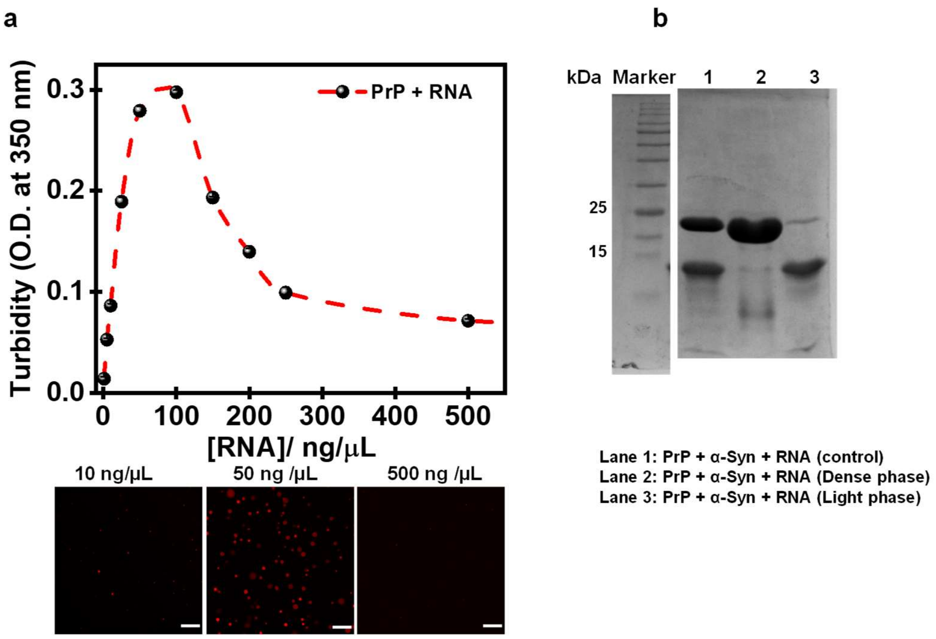


**Fig. S4 a** Solution turbidity plot for PrP as a function of polyU RNA concentration. Confocal images for Alexa-594-labeled PrP at different RNA concentrations as indicated. Scale bar: 10 μm. **b** SDS-PAGE analysis of PrP-α-Syn droplets in the presence of RNA (150 ng) using sedimentation assay.

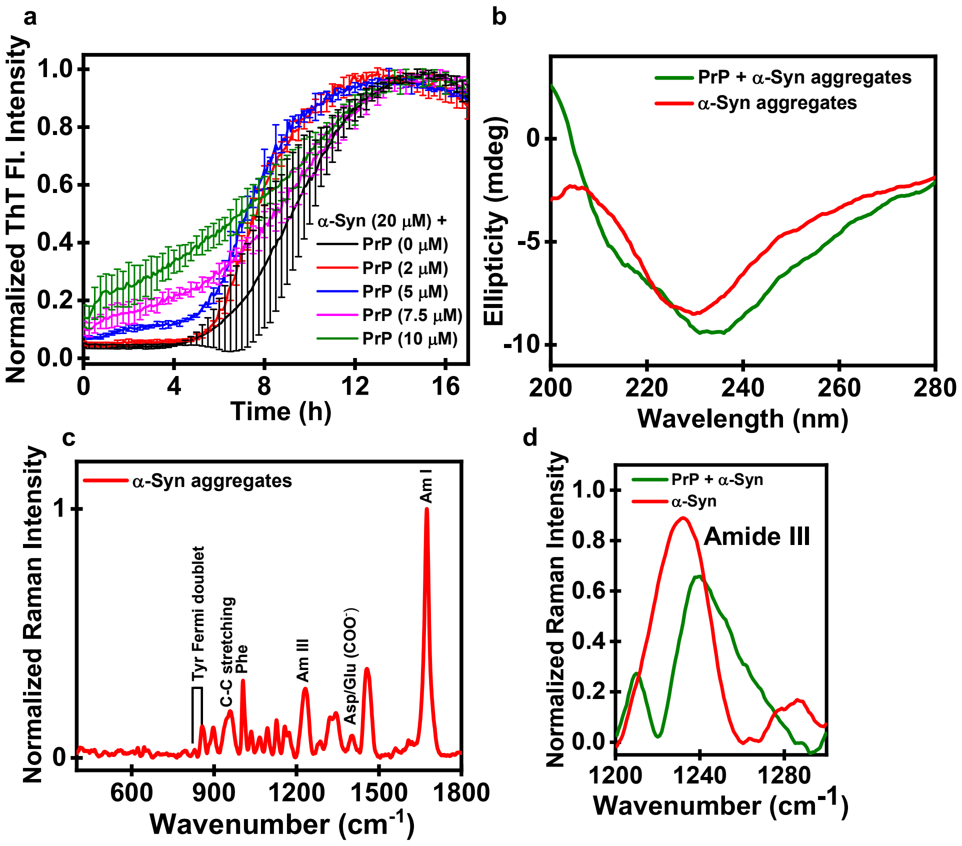


**Fig. S5** **a** ThT aggregation kinetics for α-Syn in the presence of increasing PrP concentrations indicating shortening of lag-phase. **b** Far-UV CD spectrum indicating β-rich structures for α-Syn homotypic aggregates whereas a mixture of α-helical and β-rich structure for PrP-α-Syn co-aggregates. **c** Vibrational Raman spectra of α-Syn aggregates. **d** The amide III region is shown for comparison between PrP-α-Syn co-aggregates and α-Syn homotypic aggregates.

**Table S1:** Primer sequences.

| Construct | Primer | Sequence 5'-3' |
|---|---|---|
| **α-Syn (103 Stop)** | Forward | GGACCAGTTGGGCAAGAATTAAGAAGGATGCCCACAGG |
| | Reverse | GTGGGCATCCTTCTTAATTCTTGCCCAACTGGTCC |
| **α-Syn (133 Stop)** | Forward | CCTTCTGAGGAAGGGTATTAAGACTATGAACCTGAAGCC |
| | Reverse | GGCTTCAGGTTCATAGTCTTAATACCCTTCCTCAGAAGGC |

**Table S2:** Recovered parameters from the time-resolved fluorescence anisotropy decay analysis for α-Syn labeled by IAEDANS at Cys 124 in dispersed phase and droplets.

| **α-Syn** | | **$\phi_1$ (ns) ($\beta_1$)** | **$\phi_2$ (ns) ($\beta_2$)** | **$\phi_3$ (ns) ($\beta_3$)** |
|---|---|---|---|---|
| **Residue 124** | Dispersed monomer | 0.51 ± 0.065 (0.64 ± 0.027) | 3.38 ± 0.39 (0.36 ± 0.027) | - |
| | Droplets | 0.43 ± 0.054 (0.34 ± 0.031) | 3.77 ± 0.42 (0.36 ± 0.014) | 54.26 ± 6.02 (0.31 ± 0.021) |
